# Phosphoinositide 3-kinase regulates wild-type RAS signaling to confer resistance to KRAS inhibition

**DOI:** 10.1101/2025.06.20.660715

**Authors:** Xiangyu Ge, Jaffarguriqbal Singh, Wenxue Li, Cassandra S. Markham, Christian Felipe Ruiz, Moitrayee Bhattacharyya, Yansheng Liu, Mandar Deepak Muzumdar

**Affiliations:** Yale Cancer Biology Institute, Yale University; West Haven, CT 06516, USA; Department of Pathology, Yale University School of Medicine; New Haven, CT 06510, USA; Translational Molecular Medicine, Pharmacology, and Physiology Program, Yale University; New Haven, CT 06510, USA; Department of Genetics, Yale University School of Medicine; New Haven, CT 06520, USA; Department of Pharmacology, Yale University School of Medicine; New Haven, CT 06520, USA; Molecular Cell Biology, Genetics, and Development Program, Yale University; New Haven, CT 06510, USA; Department of Biomedical Informatics & Data Science, Yale University School of Medicine; New Haven, CT 06510, USA; Department of Internal Medicine, Section of Medical Oncology, Yale University School of Medicine; New Haven, CT 06510, USA; Yale Cancer Center and Smilow Cancer Hospital, Yale University, New Haven, CT, USA

**Keywords:** KRAS, phosphoinositide 3-kinase, mitogen-activated protein kinase, pancreatic cancer, proximity protein labeling, CRISPR

## Abstract

Despite the availability of RAS inhibitors and the dependence of >90% of pancreatic ductal adenocarcinomas (PDAC) on oncogenic *KRAS* mutations, resistance to KRAS inhibition remains a serious obstacle. We show here that phosphoinositide 3-kinase (PI3K) plays a major role in this resistance through upstream activation of wild-type RAS signaling – beyond its known KRAS effector function. Combining proximity labeling, CRISPR screens, live-cell imaging, and functional assays we found that PI3K orchestrates phosphoinositide-mediated GAB1 recruitment to the plasma membrane, nucleating assembly of RAS signaling complexes that activate mitogen-activated protein kinase (MAPK) in an EGFR/SHP2/SOS1-dependent manner. We further demonstrate that inhibiting PI3K enhances sensitivity to mutant-specific KRAS inhibitors in PDAC cells, including cells with clinically identified *PIK3CA* mutations. Our findings refine RAS-PI3K signaling paradigms, reveal that PI3K-driven wild-type RAS activation drives resistance to KRAS inhibition, and illuminate new avenues for augmenting KRAS-targeted therapies in PDAC.

## INTRODUCTION

KRAS is a small GTPase that is frequently mutated in the most lethal cancers, including pancreatic ductal adenocarcinoma (PDAC), where activating mutations are present in over 90% of cases^1^. Oncogenic mutations in *KRAS* impair GTP hydrolysis resulting in elevated steady-state levels of active GTP-bound KRAS, which drives pro-tumorigenic signaling through downstream effector pathways – including the mitogen-activated protein kinase (MAPK) and phosphoinositide 3-kinase (PI3K) cascades^2^. Recent advances in the development of KRAS inhibitors have raised hopes for targeting KRAS-driven tumors, including PDAC^3,4^. However, previous preclinical studies – including our own – demonstrated that KRAS dependency is heterogeneous across *KRAS* mutant models^5,6^, and a subset of PDAC cells can even survive CRISPR/Cas9-mediated complete KRAS ablation^7^. Consistent with these findings, clinical efficacy of emerging mutant-specific KRAS inhibitors has been limited due to the rapid emergence of resistance^8–12^. Genetic and non-genetic (adaptive) resistance in multiple KRAS-driven tumor contexts largely converge on compensatory activation of receptor tyrosine kinase (RTK), RAS, and MAPK pathway effectors^11–15^, arguing that resistant cells remain addicted to RAS–MAPK signaling. Yet, how these pathways are engaged in the absence of resistance mutations to sustain tumor cell viability following KRAS inhibition remains incompletely understood.

Beyond RAS-MAPK, several studies have shown that activation of PI3K signaling may confer resistance to mutant KRAS or MAPK pathway inhibition in *KRAS* mutant cancers^16–18^. These include genetic alterations, such as activating mutations in the PI3K catalytic subunit p110α and loss-of-function mutations in the phosphatidylinositol-(3,4,5)-triphosphate (PIP_3_) phosphatase PTEN^19^. PI3K canonically activates the AKT kinases, which phosphorylate diverse substrates that regulate survival, cell proliferation, and metabolism^20^. Our previous work demonstrated that catalytic pan-class IA PI3K inhibitors not only inhibited the activation of AKT but also effectively blocked MAPK signaling in *KRAS* mutant PDAC cells subject to *KRAS* knockout^7^. This observation aligns with other prior studies reporting that PI3K inhibition reduces phosphorylated MAPK/ERK (pERK) in basal and RTK ligand-stimulated conditions in various cell lines^21–24^. These data indicate that PI3K may not be solely a downstream effector of RAS but could also play a direct regulatory role in MAPK signaling. Indeed, constitutive activation of PI3K can induce RAS-dependent MAPK signaling in some but not all cellular contexts^25,26^. Importantly, most prior studies used older, less-specific PI3K inhibitors (*e.g.*, wortmannin) and focused on ligand-stimulated signaling^24,27^. Therefore, the extent to which PI3K orchestrates wild-type RAS–MAPK signaling and the underlying mechanisms that govern this function remain unclear. Moreover, how this regulatory axis is co-opted by PDAC cells to enable resistance to KRAS inhibition has not been established.

In this study, we demonstrate that PI3K activity broadly governs RAS–MAPK signaling in basal and KRAS-inhibited states in cells. Specifically, PI3K activation enhances wild-type – but not mutant – RAS activity and MAPK phosphorylation, while PI3K inhibition reduces them. Live cell-imaging, proximity labeling proteomics, and functional experiments reveal that PI3K – through its catalytic lipid kinase function – orchestrates the membrane recruitment of the scaffold protein GAB1 to facilitate wild-type RAS activation via EGFR/SHP2/SOS1-dependent signaling complexes. These findings redefine the conventional hierarchy of RAS signaling, argue that wild-type RAS may be an effector of PI3K, and provide new mechanistic insights into how the PI3K-driven lipid environment may facilitate the assembly of pro-survival signaling hubs under therapeutic KRAS blockade. Notably, combining PI3K inhibitors with KRAS mutant-specific inhibitors elicits a synergistic effect on the viability of PDAC cells, highlighting the therapeutic potential of targeting PI3K-dependent wild-type RAS activation as a promising strategy to overcome resistance to KRAS inhibitors in PDAC.

## RESULTS

### PI3K regulates MAPK signaling in RAS wild-type cells

To test the impact of PI3K inhibition on MAPK signaling, we treated a series of *RAS* wild-type cells with the pan-class IA PI3K catalytic inhibitors GDC-0941 and ZSTK474. Specifically, we used mouse embryonic fibroblasts (MEFs) and 293 human embryonic kidney (293HEK) cells as widely used non-cancerous cell models along with *KRAS*-deficient PDAC cell lines (8988T-KO, KP4-KO) we generated by CRISPR/Cas-mediated *KRAS* knockout as cancer models of resistance to KRAS inhibition^7,28^. Across all cell lines, we observed a significant rapid reduction in pERK levels with maximal suppression achieved within 30 minutes of treatment (**Figures 1A, 1B, S1A, and S1B**). To confirm the temporal dynamics of ERK activity following PI3K inhibition using an orthologous method, we performed live-cell imaging using an ERK kinase translocation reporter (ERK-KTR)^29^ and validated the capacity of the reporter to serve as a surrogate for pERK levels (**Figures S1C and S1D**). PI3K inhibition rapidly attenuated ERK activity within minutes in *KRAS*-deficient PDAC and 293HEK cells (**Figure S1E**). We further observed that catalytic inhibitors targeting specific PI3K p110 isoforms (alpelisib (BYL-719) for p110α, TGX-211 for p110β, idelalisib for p110δ) also reduced pERK levels to varying degrees (**Figure 1C**). Strikingly, across all cell lines, pERK suppression correlated with pAKT inhibition with maximal suppression achieved by pan-PI3K isoform blockade (**Figure 1D**), indicating functional redundancy of class IA p110 isoforms in regulating MAPK signaling.

**Figure 1.**
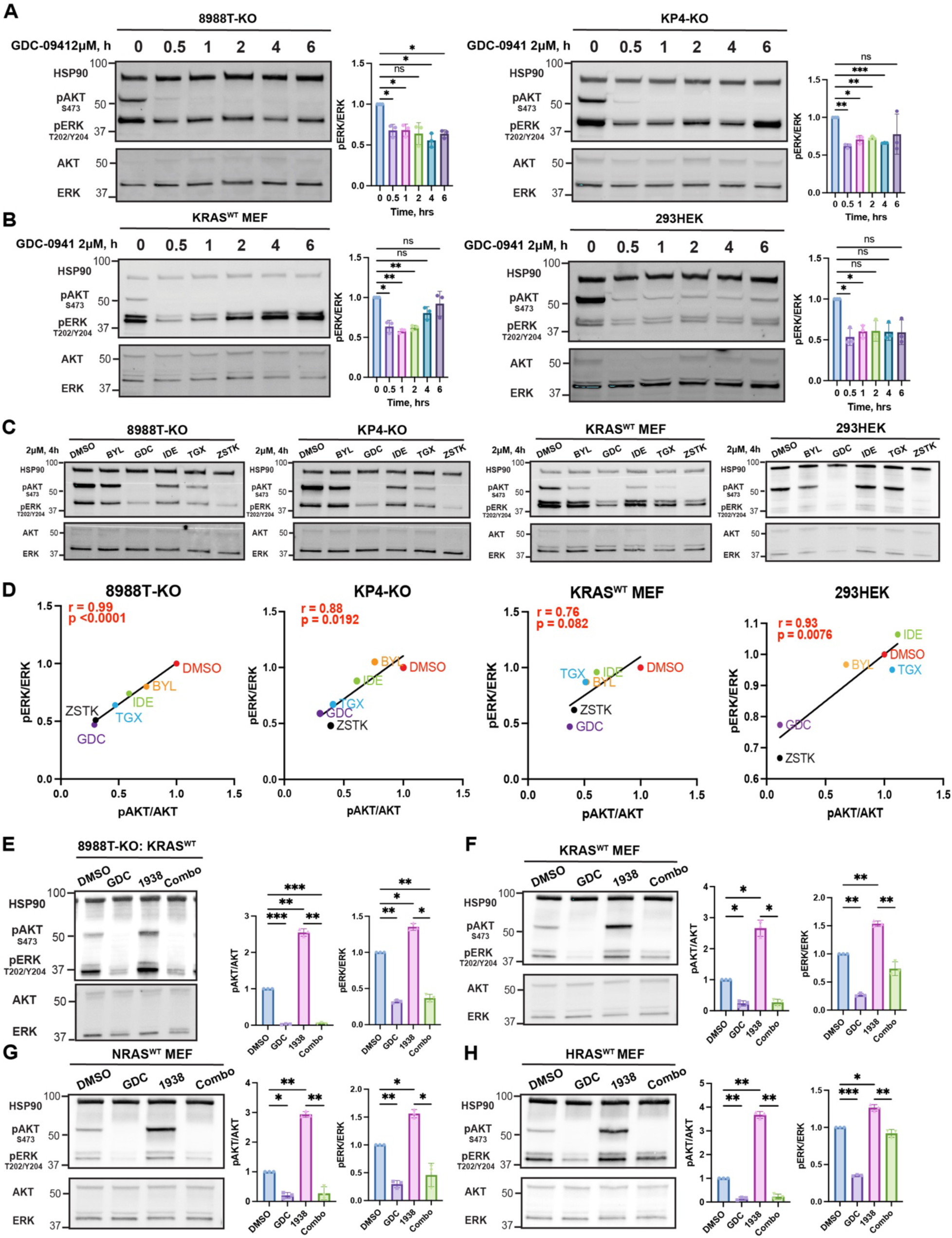
PI3K regulates MAPK signaling in wild-type RAS cells. (A) Representative immunoblots showing the effect of PI3K inhibition (GDC-0941, 2 μM) over a 6-hour time course on phosphorylated ERK1/2 (pERK^T202/Y204^) and phosphorylated AKT (pAKT^S473^) in *KRAS*-deficient PDAC clones (8988T-KO, KP4-KO). HSP90 is loading control. Bar graphs quantify pERK/ERK and pAKT/AKT levels normalized to time 0 (mean ± SD, n = 3 biological replicates). ns = not significant, * p < 0.05, ** p < 0.01 *** p < 0.001, repeated measures ANOVA (rmANOVA) with Dunnett’s post-hoc test. (B) Representative immunoblots showing the effect of PI3K inhibition (GDC-0941, 2 μM) over a 6-hour time course on pERK1/2 and pAKT in non-malignant cells (293HEK, KRAS^WT^ MEF cells). Bar graphs quantify pERK/ERK and pAKT/AKT levels normalized to time 0 (mean ± SD, n = 3 biological replicates). ns = not significant, * p < 0.05, ** p < 0.01, repeated measures ANOVA (rmANOVA) with Dunnett’s post-hoc test. (C) Immunoblots showing pERK1/2 and pAKT levels in designated cell lines following 4-hour treatment with isoform-selective PI3K inhibitors (2 μM each): BYL719 (BYL, p110α), GDC-0941 (GDC, pan-class IA), idelalisib (IDE, p110δ), TGX221 (TGX, p110β), and ZSTK474 (ZSTK, pan-class IA). HSP90 is loading control. (D) Correlation between pAKT/AKT and pERK/ERK (normalized to DMSO) across inhibitor treatments was assessed in each cell line in (**C**), indicating the extent to which PI3K signaling activity (pAKT) is associated with MAPK output (pERK). Pearson correlation coefficients (r) and associated p-values are shown. (E-H) Representative immunoblots and quantification of pERK1/2 and pAKT following treatment with DMSO (solvent), 2 μM GDC-0941 (GDC), 1 μM PI3K activator 1938, or their combination for 0.5 hour in 8988T-KO (**E**), KRAS^WT^ MEF (**F**), NRAS^WT^ MEF (**G**), and HRAS^WT^ MEF (**H**) cells. Bar graphs represent mean ± SD normalized to DMSO from n = 3 biological replicates. * p < 0.05, ** p < 0.01, *** p < 0.001, repeated measures ANOVA (rmANOVA) with Šidák’s post-hoc test.

We next evaluated whether PI3K activation would conversely induce MAPK signaling in *RAS* wild-type cells. To accomplish this, we took advantage of a recently developed allosteric PI3Kα activator UCL-TRO-1938 (1938)^30^. As expected, 1938 increased pAKT levels, and this effect was abolished by a catalytic PI3Kα inhibitor in 293HEK and *KRAS*-deficient PDAC cells (**Figure S1F**). Remarkably, 1938 treatment increased pERK levels in 8988T-KO PDAC cells reconstituted with KRAS^WT^ by lentiviral transduction (8988T-KO: KRAS^WT^), which was abrogated by co-treatment with GDC-0941 (**Figure 1E)**. These results were recapitulated across *Ras*-less MEFs reconstituted to express a single Ras isoform (Kras, Hras, or Nras)^31^, suggesting that PI3K-mediated bidirectional regulation of MAPK signaling extends across wild-type RAS isoforms (**Figures 1F-1H)**. Importantly, these effects occurred in the absence of exogenous RTK ligand treatment. Collectively, these findings establish PI3K as a regulator of MAPK signaling across varied non-cancerous and cancerous cells, including in the mutant KRAS-inhibited state. Furthermore, this regulation occurs irrespective of RAS or p110 isoform.

### PI3K regulates wild-type RAS activity

We next delineated whether PI3K exerted its effects on MAPK signaling by altering RAS activity. RAS-GTP ELISA assays demonstrated PI3K inhibition decreased RAS-GTP levels in 8988T-KO: KRAS^WT^ PDAC cells, whereas PI3K activation increased levels (**Figure 2A**). Similarly, MEFs expressing single Ras isoforms exhibited reduced RAS-GTP levels with PI3K inhibitor treatment, supporting a tight link between PI3K and basal RAS activity (**Figure 2B**). We confirmed this effect using two orthogonal approaches: 1) active RAS pull-down using the RAS-binding domain (RBD) of CRAF (**Figure 2C**); and 2) reciprocally probing for interactions of CRAF and BRAF with KRAS by green fluorescent protein (GFP)-Trap immunoprecipitation of GFP-tagged KRAS^WT^ in reconstituted 8988T-KO cells (8988T-KO: GFP-KRAS^WT^) (**Figure 2D**). In contrast to KRAS^WT^, PI3K inhibition had no impact on RAS-GTP levels in mutant KRAS-expressing cells (**Figures 2E and 2F**). As a consequence, KRAS^G12V^ expression abolished the effect of PI3K inhibition on ERK activity in 293HEK cells assessed by both immunoblotting and ERK-KTR-based imaging (**Figures S1C and S1E**). Furthermore, mutant KRAS expression reduced the impact of PI3K inhibition on pERK levels and decreased sensitivity to PI3K inhibition in PDAC cells, irrespective of *KRAS* mutant variant (**Figures S2A and S2B**). We reasoned that if PI3K functions at the level of RAS itself, upstream activation of wild-type RAS by RTKs should be sensitive to PI3K inhibition. Indeed, EGF-stimulated activation of MAPK signaling was markedly reduced by concurrent PI3K inhibition in RAS^WT^ cells (**Figure S2C**). In contrast, we expected that constitutive activation of the immediate downstream RAS effector BRAF should abolish the impact of PI3K inhibition on MAPK signaling. As predicted, pharmacologic modulation of PI3K activity had no effect on pERK levels in *Ras*-less MEFs expressing BRAF^V600E^ (**Figure S2D)**, mirroring observations in MEFs expressing KRAS^G12V^ (**Figure 2G**). We confirmed this finding in the *KRAS^WT^*; *BRAF^V600E^*mutant PDAC cell line BxPC3 (**Figure S2E**). Together, these data reveal a non-canonical role for PI3K as an upstream regulator of wild-type RAS activity.

**Figure 2.**
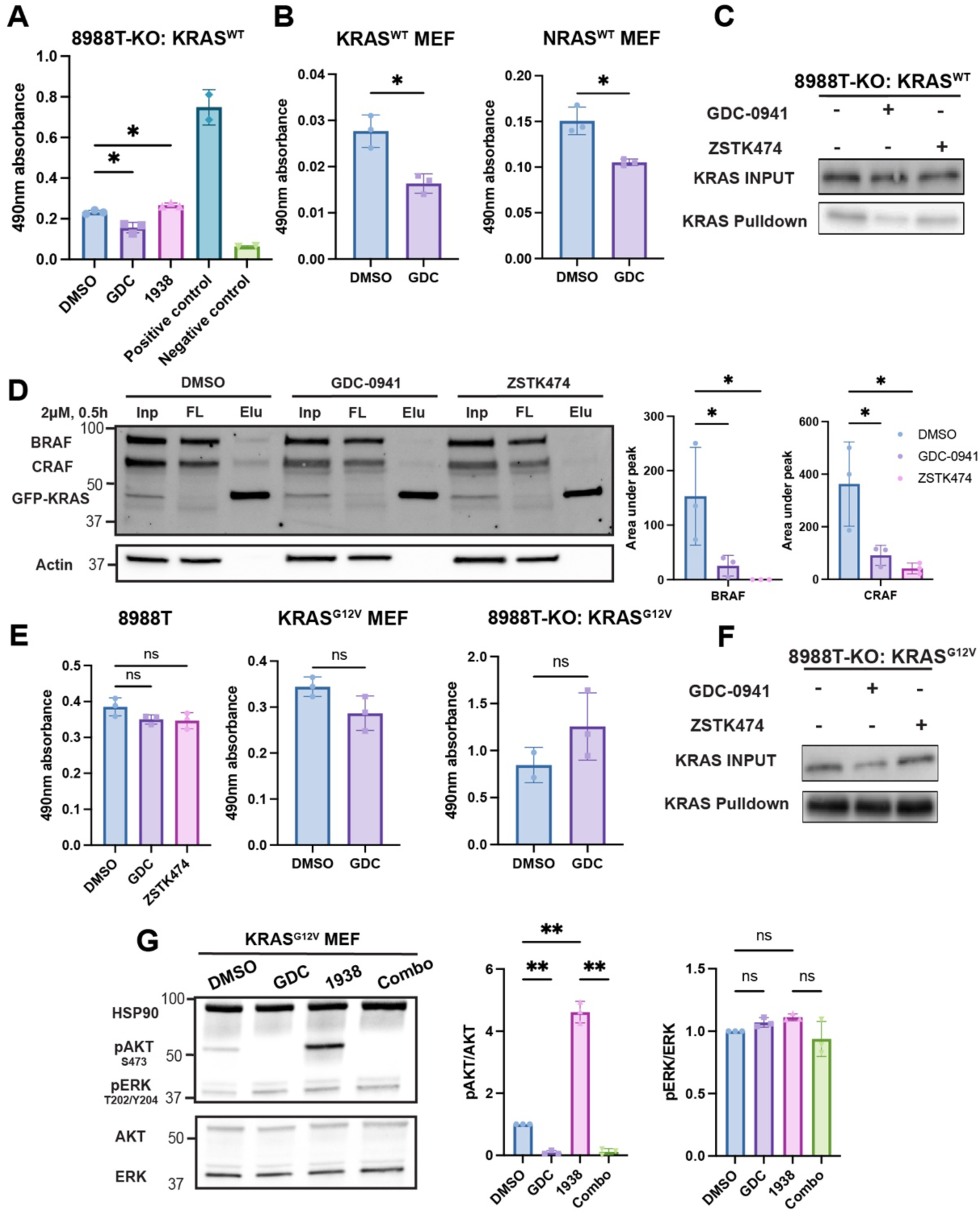
PI3K regulates wild-type but not mutant RAS activity. (A) G-LISA RAS activation assays in *KRAS*-deficient PDAC cells reconstituted with wild-type KRAS (8988T-KO: KRAS^WT^) and treated with GDC-0941 (GDC, 2 μM), 1938 (1 μM), or solvent (DMSO) control for 30 minutes RAS-GTP levels are measured as absorbance (mean ± SD, n = 3 biological replicates per condition). Positive (active RAS control protein) and negative (buffer only) controls are shown. * p < 0.05, one-way ANOVA with Tukey’s post-hoc test. (B) G-LISA RAS activation assays in single RAS-expressing (KRAS^WT^, NRAS^WT^) MEFs treated with GDC. RAS-GTP levels are measured as absorbance (mean ± SD, n = 3 biological replicates per condition). * p < 0.05, paired two-tailed t-test. (C) Representative immunoblots of RAS-GTP (pulldown assay using the RAF-RBD) and input lysate in 8988T-KO: KRAS^WT^ cells upon PI3K inhibitors treatment (2 μM, 4 hours). (D) Representative immunoblots of GFP-tagged KRAS (GFP-KRAS) and associated effectors (BRAF and CRAF) following GFP nano-trap pulldown of KRAS assays upon PI3K inhibition (GDC-0941 or ZSTK474, 2 μM) in 8988T-KO cells reconstituted with GFP-KRAS (8988T-KO: GFP-KRAS^WT^ cells. Densitometric quantification (mean ± SD, n = 3 biological replicates). *p < 0.05, one-way ANOVA with Tukey’s post hoc test. (E) G-LISA RAS activation assays in mutant KRAS-expressing cells (8988T parental cells, single RAS-expressing KRAS^G12V^ MEFs, and 8988T-KO cells reconstituted with KRAS^G12V^ (8988T-KO: KRAS^G12V^)) treated with GDC-0941 (GDC, 2 μM), ZSTK474 (2 μM), or solvent (DMSO) control for 4 hours. RAS-GTP levels are measured as absorbance (mean ± SD, n = 3 biological replicates per condition). ns = non-significant, one-way ANOVA with Tukey’s post-hoc test or paired two-tailed t-test. (F) Representative immunoblot of RAS-GTP pulldown assay using the RAF-RBD in 8988T-KO: KRAS^G12V^ cells upon PI3K inhibitors treatment (2 μM, 4 hours). (G) Representative immunoblots and quantification of phosphorylated ERK1/2 (pERK^T202/Y204^) and phosphorylated AKT (pAKT^S473^) following treatment with DMSO (solvent), 2 μM GDC-0941 (GDC), 1 μM PI3K activator 1938, or their combination for 0.5 hour in KRAS^G12V^ MEFs. HSP90 is loading control. Bar graphs represent mean ± SD normalized to DMSO from n = 3 biological replicates. ns = non-significant, ** p < 0.01, repeated measures ANOVA (rmANOVA) with Šidák’s post-hoc test.

### PI3K-mediated wild-type RAS signaling is altered by PTEN but not AKT

Class I PI3Ks phosphorylate phosphatidylinositol-(4,5)-bisphosphate (PI(4,5)P_2_), resulting in the generation of PIP_3_^32^, which is dephosphorylated by PTEN^19^. As the effect of 1938 on pERK levels was abolished by catalytic PI3K inhibition (**Figures 1E-H**), we reasoned that PIP_3_ likely mediates MAPK signaling. To substantiate this hypothesis, we knocked down PTEN in 8988T-KO: KRAS^WT^ cells, which in steady-state conditions increased pAKT but not pERK levels. This may be attributable to compensatory enrichment of PI(3,4)P_2_ at the membrane, which – like PIP_3_ – also binds the pleckstrin homology (PH) domain of AKT^33,34^. Importantly, PTEN knockdown further enhanced 1938-induced pERK and pAKT levels **(Figure 3A**), supporting sustained membrane PIP_3_ in amplifying MAPK signaling output. To determine whether PI3K regulates wild-type RAS signaling through its canonical effector AKT, we treated PDAC and MEF cells expressing KRAS^WT^ with the allosteric AKT inhibitor MK-2206^35^. Notably, MK-2206 treatment did not reduce basal pERK levels nor suppress the 1938-induced increase in pERK observed in KRAS^WT^-expressing 8988T-KO and MEF cells (**Figure 3B)**. Beyond AKT, PIP_3_ recruits diverse proteins to the plasma membrane including GTPase regulators^36–38^, kinases^39,40^, adaptor/scaffold proteins^41^ and phosphatases^42^. Wild-type RAS proteins are tightly regulated by guanine nucleotide exchange factors (GEFs) and GTPase-activating proteins (GAPs), including some that harbor PH domains capable of binding PIP_3_^43^. To test whether PI3K regulates GEFs or GAPs to modulate RAS activity, we perturbed the RAS regulatory network by overexpressing the RAS-GEF *SOS1* or knocking down the RAS-GAP *NF1*. Both *SOS1* overexpression and *NF1* knockdown increased basal pERK levels, but pERK was still effectively suppressed by PI3K inhibition (**Figures S3A and S3B**). Knockdown of additional RAS-GAPs (*RASA1*, *RASAL1*) also did not impact the effect of PI3K inhibition on pERK (**Figure S3C**). Together, these findings indicate that PI3K sustains basal MAPK signaling independently of AKT and canonical RAS-GEFs and GAPs, arguing that additional PIP_3_-associated proteins may mediate the impact of PI3K on RAS–MAPK signaling.

**Figure 3.**
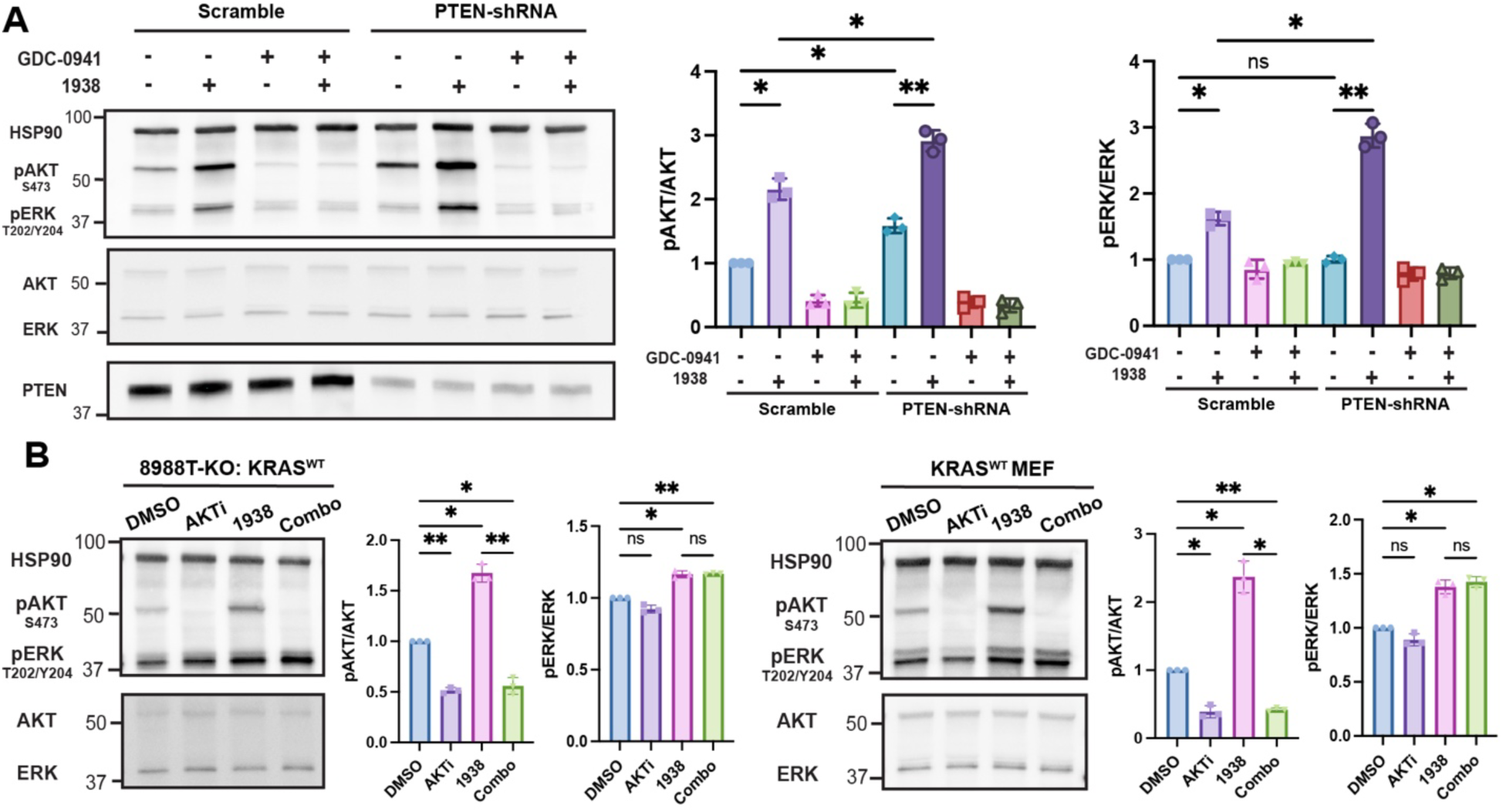
PI3K-mediated RAS wild-type signaling is altered by PTEN but not AKT. (A) Representative immunoblots of phosphorylated ERK1/2 (pERK^T202/Y204^) and phosphorylated AKT (pAKT^S473^) in *PTEN*-intact (Scramble) and *PTEN*-knockdown (PTEN-shRNA) 8988T-KO: KRAS^WT^ cells treated with DMSO, 1938 (1 μM), GDC-0941 (2 μM), or the combination for 15 minutes (peak response of pAKT levels with 1938 treatment). HSP90 is loading control. Bar graphs show quantification of pAKT/AKT and pERK/ERK normalized to DMSO (mean ± SD, n = 3 biological replicates). ns = non-significant, * p < 0.05, ** p < 0.01, repeated measures ANOVA (rmANOVA) with Šidák’s post-hoc test. (B) Representative immunoblots and quantification of pERK1/2 and pAKT following treatment with DMSO (solvent), 2 μM MK-2206 (AKTi), 1 μM PI3K activator 1938, or their combination (Combo) for 0.5 hour in 8988T-KO: KRAS^WT^ and single RAS-expressing KRAS^WT^ MEF cells. Bar graphs represent mean ± SD normalized to DMSO from n = 3 biological replicates. ns = non-significant, * p < 0.05, ** p < 0.01, repeated measures ANOVA (rmANOVA) with Šidák’s post-hoc test.

### Protein proximity labeling defines the PI3K-dependent KRAS interactome

To identify PI3K-dependent KRAS regulators in an unbiased manner, we employed a proximity-based protein labeling strategy using the BioID system^44^. We generated human PDAC cell lines expressing BirA-tagged KRAS^WT^, oncogenic KRAS (KRAS^G12V^), or two localization controls: a cytosol-retained KRAS mutant (KRAS^C185S^) and the membrane-localized KRAS hypervariable region (KRAS^HVR^) (**Figure S4A**). By introducing these vectors into *KRAS*-deficient PDAC cells (8988T-KO), we sought to eliminate interference from endogenously expressed KRAS. Following treatment with biotin with or without the PI3K inhibitor GDC-0941 for 16 hours, proteins biotinylated by BirA were enriched using streptavidin affinity purification and subjected to data-independent acquisition mass spectrometry (DIA-MS) analysis^45^ (**Figure 4A and Table S1**). As expected, canonical RAS effectors (ARAF, BRAF, and CRAF (RAF1)) were enriched in KRAS^G12V^ mutant PDAC cells under basal conditions, relative to KRAS^WT^ and localization controls, highlighting the importance of a GTP-bound active conformation for effective effector engagement (**Figure 4B**). Furthermore, the basal KRAS^HVR^ interactome enriched for membrane-associated proteins (*e.g.,* EGFR, GAB1/2, SHP2 (PTPN11)) but, importantly, not RAF proteins, which bind the RAS G-domain (**Figure 4B**).

**Figure 4.**
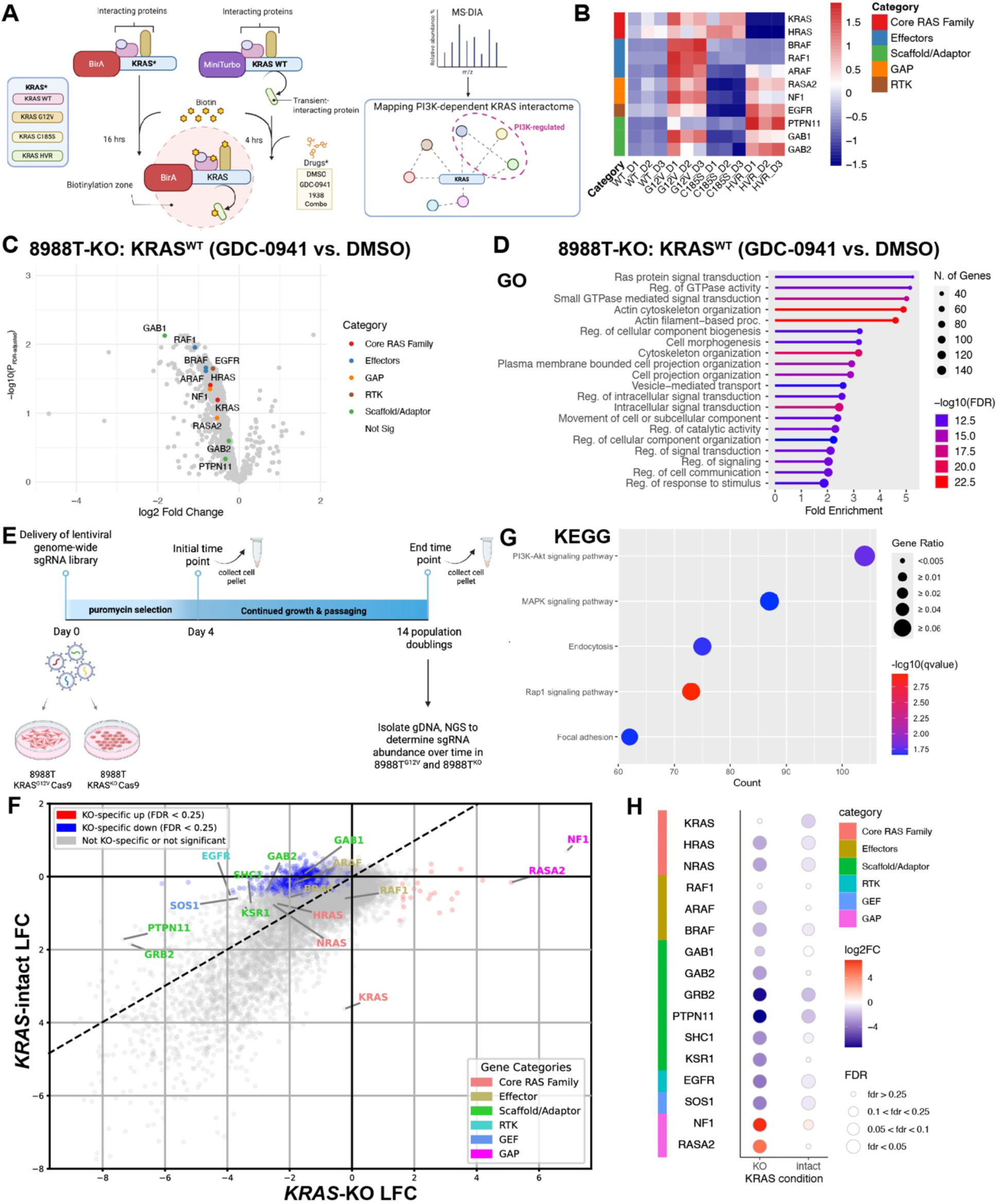
Protein proximity labeling identifies PI3K-dependent KRAS interactors. (A) Schematic illustrating protein proximity labeling workflow. *KRAS* deficient PDAC cells reconstituted with various KRAS variants were treated with biotin and pharmacologic modulators of PI3K activity and protein lysates were subject to streptavidin pull-down prior to data-independent acquisition mass spectrometry (DIA-MS). (B) Heatmap showing row-normalized relative abundance of core RAS family proteins, effectors, scaffolds/adaptors, GAPs, and RTKs in detected in proximity to KRAS in reconstituted 8988T-KO cells in basal conditions. (C) Volcano plot (log2 fold-change (FC) vs. −log10(adjusted p-value) of 8988T-KO: KRAS^WT^ cells treated with GDC-0941 vs DMSO showing reduced association of GAB1 and multiple scaffold/adaptor proteins upon PI3K inhibition. (D) Gene Ontology (GO) enrichment analysis of proteins significantly decreased following GDC-0941 treatment reveals enrichment of RAS-related signaling, actin cytoskeleton, and membrane-associated functions. (E) Schematic of genome-scale loss-of-function CRISPR screen (Brunello sgRNA library) performed in pools of 8988T *KRAS* intact and KO clones. Cells were grown for 14 population doublings following selection and genomic DNA was sequenced for sgRNA abundance at endpoint versus initial point. (F) Scatterplot showing the log2 fold-change (LFC) in abundance between end and initial time points of all genes in 8988T *KRAS*-intact and KO cells (calculated using MAGeCK). Blue dots indicate *KRAS* KO-specific dependencies (LFC < –1 and FDR < 0.25), while red dots indicate genes whose knockout confers a *KRAS* KO-specific growth advantage (LFC > 1 and FDR < 0.25). Dotted line separates genes associated with a >1 difference in LFC between *KRAS*-intact and KO cells with greater dependence in KO cells to the left of the line. RAS– MAPK signaling regulators identified in proximity biotinylation experiments in (**C**) are labelled. (G) KEGG pathway enrichment analysis was performed on genes that were significantly and specifically depleted in *KRAS*-KO cells (but not in *KRAS*-intact cells) in the genome-wide CRISPR screen. These *KRAS* KO-specific dependencies are enriched for pathways including PI3K signaling, MAPK signaling, and focal adhesion, highlighting key signaling and structural pathways required for cell survival in the absence of *KRAS*. (H) Dot plots showing relative changes in sgRNA abundance and FDR for key RAS family proteins, effectors, scaffolds/adaptors, GAPs, and RTKs in *KRAS*-intact and KO cells from the genome-wide CRISPR screen. Notably, positive regulators of MAPK signaling were more depleted in *KRAS*-KO cells while negative regulators (*e.g.,* GAPs) were more enriched.

PI3K inhibition led to significantly fewer proteins interacting with KRAS^WT^ than in control (DMSO) conditions (**Figure 4C and Table S2**). Functional overrepresentation analyses revealed that these proteins are involved in cytoskeletal organization and intracellular signaling, especially phosphoinositide and RAS–MAPK signaling-related proteins (**Figures 4D and S4B**). Specifically, PI3K inhibition led to loss of interaction of KRAS with core regulatory components of the MAPK signaling pathway, including other RAS family members (HRAS, NRAS), effectors (ARAF, BRAF, RAF1), GAPs (NF1, RASA2), RTKs (EGFR), and scaffold proteins (GAB1/2, SHP2) (**Figure 4C**). Importantly, these changes in the interactome (with the notable exception of RAF proteins) were also found in KRAS^HVR^-expressing cells, but not in KRAS^C185S^-expressing cells, treated with the PI3K inhibitor (**Figure S4C and Table S2**), suggesting that these interactions occur at the membrane. Notably, these interactions were not significantly altered in mutant KRAS^G12V^-expressing cells (**Figure S4D and Table S2**), consistent with PI3K having a greater regulatory role on KRAS^WT^ (**Figures 2 and S2**). Moreover, we confirmed the concordant impact of PI3K inhibition on the KRAS^WT^ interactome in 293HEK cells (**Figures S4A and S4E and Table S2**). Collectively, these data indicate that PI3K regulates interactions of KRAS with core MAPK regulators, effectors, and scaffolds independent of cell-type or even cancer state.

### KRAS-deficient cells are dependent on the PI3K-regulated RAS interactome

We previously found that *KRAS*-deficient PDAC cells exhibited enhanced sensitivity to PI3K inhibition relative to *KRAS*-intact cells and that this was due in part to MAPK suppression, as this could be reversed by expression of constitutively active MEK (MEK-DD)^7^ or mutant KRAS (**Figure S2B**). Therefore, we hypothesized that *KRAS*-deficient cells would show greater dependency on the PI3K-regulated RAS interactome. To test this hypothesis, we performed loss-of-function CRISPR screens in pools of *KRAS*-intact and *KRAS*-deficient 8988T clones using the Brunello genome-scale lentiviral sgRNA library^46^ (**Figure 4E**). MAGeCK analyses^47^ delineated positively and negatively selected genes in *KRAS*-deficient (relative to intact) cells over >14 population doublings (**Figure 4F and Table S3**). As expected, sgRNAs targeting *KRAS* were selectively depleted in *KRAS*-intact cells (**Figure 4F**). Conversely, sgRNAs targeting *NF1* and *RASA2* – two well-established RAS-GAPs with selectivity for KRAS/HRAS and NRAS, respectively^48^ – were specifically enriched in *KRAS*-deficient cells (**Figure 4F**). *KRAS*-deficient cells further showed selective depletion of genes associated with PI3K and MAPK signaling (**Figure 4G**). Importantly, among genes encoding proteins whose interactions with KRAS^WT^ were reduced by PI3K inhibition (**Figure 4C**), 25.1% of them were also genetic dependencies specifically in *KRAS*-deficient cells, including many known RAS regulators, scaffolds, and effectors (**Figure 4H**). In contrast, only 1% were dependencies in *KRAS*-intact cells. These data suggest that PI3K regulates the interaction of RAS proteins with their essential signaling components, underlying PI3K inhibitor sensitivity in the KRAS-inhibited state.

### GAB1 nucleates PI3K-dependent KRAS–MAPK signaling

To decipher the precise mechanisms by which PI3K may promote RAS–MAPK signaling, we performed protein proximity labeling in cells treated with the PI3K activator 1938. Since 1938 functions within a short-lived effective window to regulate MAPK signaling (**Figure S5A**), we turned to MiniTurbo-ID, an optimized proximity labeling technique with significantly faster biotinylation kinetics^49^ than conventional BioID. We treated serum-starved PDAC cells stably expressing MiniTurbo-tagged KRAS^WT^ (8988T-KO: MiniTurbo-KRAS^WT^) with either vehicle control (DMSO), PI3K activator (1938), PI3K inhibitor (GDC-0941), or a combination of both drugs for four hours (**Figure S5B**) and performed DIA-MS analysis on streptavidin-enriched proteins (**Table S1**). As expected, proteins that exhibited reduced KRAS interactions in cells treated with the combination of 1938 and GDC-0941 were related to phosphoinositide and MAPK signaling (**Figure S5C and Table S2**). To identify KRAS interactors dependent on the catalytic activity of PI3K, we focused on proteins whose interactions with KRAS increased upon PI3K activation and decreased under PI3K inhibition. Thirteen proteins fit this pattern, including GTPase regulators (FGD1, RGL3, DENDD2A, ARAP2), kinases/phosphatases (CDK14, PKN3, PRKAG2, SYNJ2, MTMR12), and other molecules (SHTN1, PLEKHA2, AHNAK2, GAB1), many of which harbor PH domains for phosphoinositide binding and regulate the cytoskeleton, cell adhesion, or motility (**Figure 5A)**. Of these proteins, GAB1 exhibited the most significantly decreased interaction in cells subject to PI3K inhibition following 1938 treatment (**Figures S5D and Table S2**).

**Figure 5.**
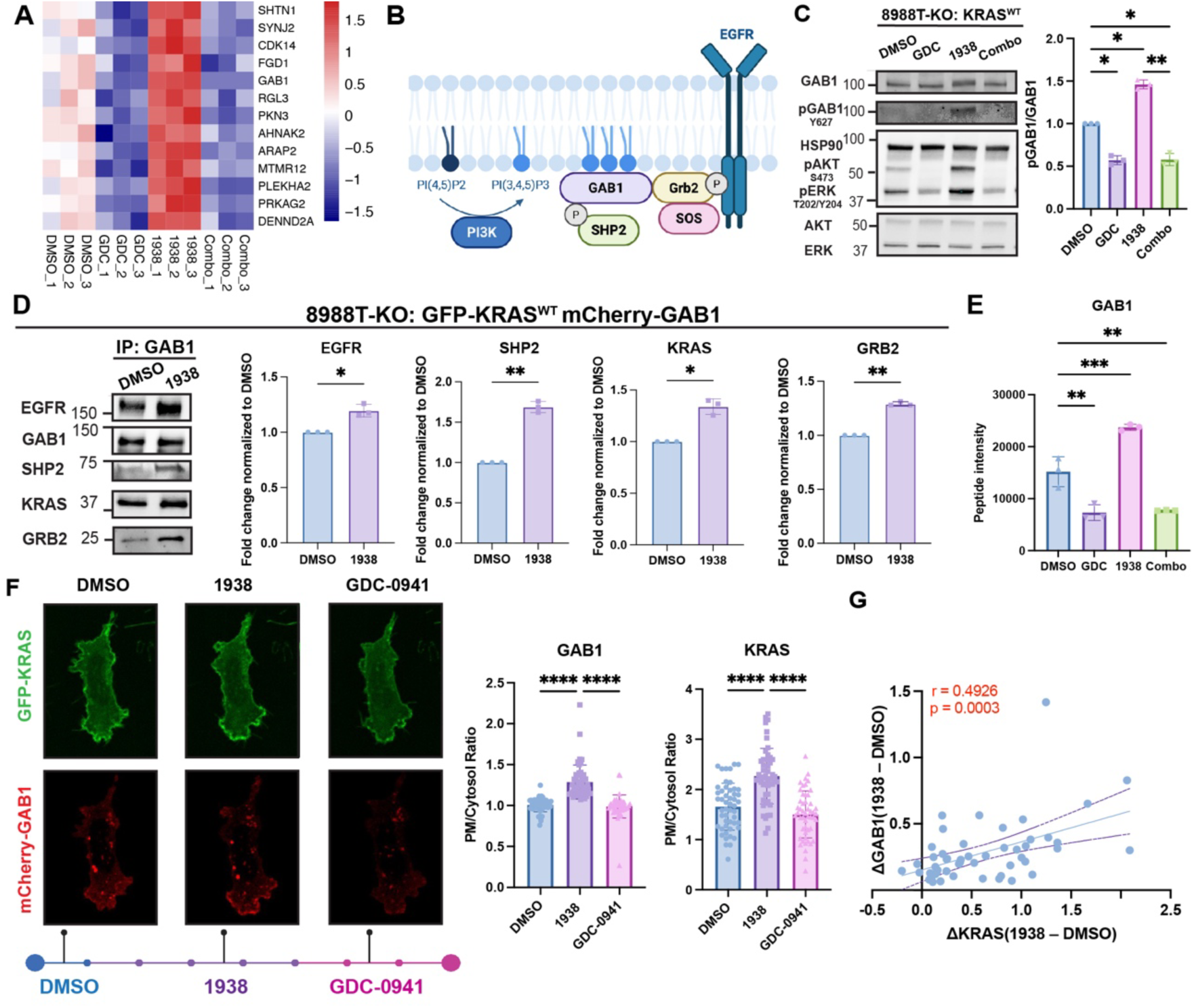
GAB1 nucleates PI3K-driven RAS–MAPK signaling. (A) Heatmap showing row-normalized relative abundance of candidate PI3K-dependent KRAS interactors based on proximity biotinylation analysis in 8988T-KO: MiniTurbo-KRAS^WT^ cells treated with DMSO solvent, GDC-0941 (GDC, 2 μM), 1938 (1 μM), or both (Combo) for 0.5 hours. Each biologic replicate (1, 2, and 3) for each condition is shown. (B) Schematic model illustrating GAB1 as a nucleating protein to initiate downstream RAS– MAPK signaling. Created with BioRender.com. (C) Representative immunoblots of phosphorylated GAB1 (pY627), phosphorylated ERK1/2 (pERK^T202/Y204^), and phosphorylated AKT (pAKT^S473^) in 8988T-KO: KRAS^WT^ cells upon treatment with PI3K inhibitor (GDC-0941, 2 μM), activator (1938, 1 μM), or both (Combo). HSP90 is loading control. Bar graphs show quantification of pGAB1/GAB1 normalized to DMSO (mean ± SD, n = 3 biological replicates). ns = non-significant, * p < 0.05, ** p < 0.01, repeated measures ANOVA (rmANOVA) with Šidák’s post-hoc test. (D) Representative immunoblots of mCherry-GAB1 interactomes from RFP-Trap immunoprecipitation of GAB1 in 8988T-KO: GFP-KRAS^WT^ cells treated with 1938 (1 μM, 0.5h). Bar graphs show quantification normalized to DMSO (mean ± SD, n = 3 biological replicates). * p < 0.05, ** p < 0.01, paired two-tailed t-test. (E) Quantification of GAB1 peptide intensities in (**A**). Data are presented as mean ± SD, n = 3 biological replicates. ** p < 0.01, *** p < 0.001, repeated measures ANOVA (rmANOVA) with Dunnett’s post-hoc test) (F) Confocal live-cell imaging of GFP-KRAS and mCherry-GAB1 in 8988T-KO: GFP-KRAS^WT^, mCherry-GAB1 cells sequentially treated with DMSO, 1938 (1 μM), and GDC-0941 (2 μM) at the time points shown. Bar graphs show quantification of plasma membrane (PM) to cytosol intensity ratio of GAB1 and KRAS (see **Methods**). Data are represented as mean ± SD from n = 50 cells from at least 10 biological replicates for each condition. **** p < 0.001, repeated measures ANOVA (rmANOVA) with Šidák’s post-hoc test. (G) Positive correlation between the change in GAB1 and KRAS PM to cytosol intensity following 1938 treatment of cells in (**F**). Pearson correlation coefficient (r) and p-value are shown.

GAB1 is a signaling scaffold that can be recruited by PIP_3_ to the plasma membrane and associates with GRB2 and SHP2 to potentiate RAS–MAPK signaling in EGF-stimulated conditions^21,50^ (**Figure 5B**). Given the selective dependence of *KRAS*-deficient cells on GAB1, GRB2, and SHP2 (**Figure 4H**), we reasoned that GAB1 may also mediate RAS–MAPK signaling in basal conditions and in the context of direct PI3K activation. Indeed, our results showed that phosphorylation of GAB1 at Y627, a well-characterized docking site for SHP2^51^, was regulated by PI3K activity, as 1938 enhanced Y627 phosphorylation while GDC-0941 abrogated it in KRAS^WT^ PDAC cells (**Figure 5C**). To validate the PI3K-dependent interaction between GAB1 and KRAS, we developed a dual-labeled PDAC cell line co-expressing mCherry-tagged GAB1 and GFP-tagged KRAS^WT^ (8988T-KO: GFP-KRAS^WT^ mCherry-GAB1). Red fluorescent protein (RFP)-Trap immunoprecipitation of mCherry-GAB1 showed increased GAB1 interactions with several known binding partners including GRB2, SHP2, and EGFR following 1938 treatment (**Figure 5D**). Notably, the association between GAB1 and KRAS also increased (**Figures 5D and 5E**), confirming our proximity biotinylation experiments. We corroborated these findings using live-cell confocal microscopy to assess the subcellular co-localization of GAB1 and KRAS at the membrane with perturbations of PI3K activity. Upon treatment with 1938, GAB1 rapidly translocated from the cytosol to the plasma membrane, likely due to its recruitment by PIP_3_, where it co-localized with KRAS in proportionally increased levels (**Figures 5F and 5G**). This effect was reversed by PI3K inhibition (**Figure 5F).** These findings position GAB1 as a PI3K-regulated nucleating scaffold that orchestrates the assembly of key signaling complexes at the plasma membrane to promote RAS activation.

We next studied how GAB1 contributes to RAS–MAPK activation in PDAC cells. Phosphorylation of GAB1 at Y627 can be induced by several RTKs^51,52^, and experiments using selective inhibitors of EGFR argue that GAB1 phosphorylation in basal and 1938-stimulated conditions was dependent on EGFR in *KRAS*-deficient PDAC cells (**Figure 6A**). Strikingly, 1938 increased EGFR phosphorylation, suggesting that PI3K-induced EGFR activation may enhance MAPK signaling. Indeed, EGFR inhibition reduced pERK levels induced by 1938 (**Figure 6A**). GAB1 phosphorylation at Y627 recruits SHP2 and GAB1 associates with GRB2, which in turn can recruit SOS1 (**Figure 5B**) to activate RAS–MAPK signaling. Consistent with these events contributing to PI3K-mediated MAPK signaling, both SHP2 and SOS1 inhibition reduced pERK levels in basal conditions and abolished the effect of 1938 on pERK (**Figures 6B and 6C**). Importantly, EGFR, GRB2, SHP2, and SOS1 were all selective dependencies in *KRAS*-deficient cells (**Figure 4H**), supporting their importance in maintaining RAS–MAPK signaling in the KRAS-inhibited state. A recent study reported that mutant PI3K-AKT signaling is spatially organized at focal adhesions and is dependent on FAK activity^53^. To test whether wild-type PI3K-dependent RAS–MAPK signaling is also FAK-dependent, we treated 8988T-KO: KRAS^WT^ cells with the FAK inhibitor defactinib. Although defactinib partially reduced pAKT levels (relative to PI3K inhibition), pERK levels were not significantly impacted in basal nor 1938-stimulated conditions (**Figure 6D**), arguing that FAK-dependent PI3K activity compartmentalized at focal adhesions^53^ does not primarily mediate RAS–MAPK signaling. Instead, PI3K orchestrates the assembly of GAB1-nucleated EGFR/GRB2/SHP2/SOS1 signaling complexes that drive RAS– MAPK pathway activation.

**Figure 6.**
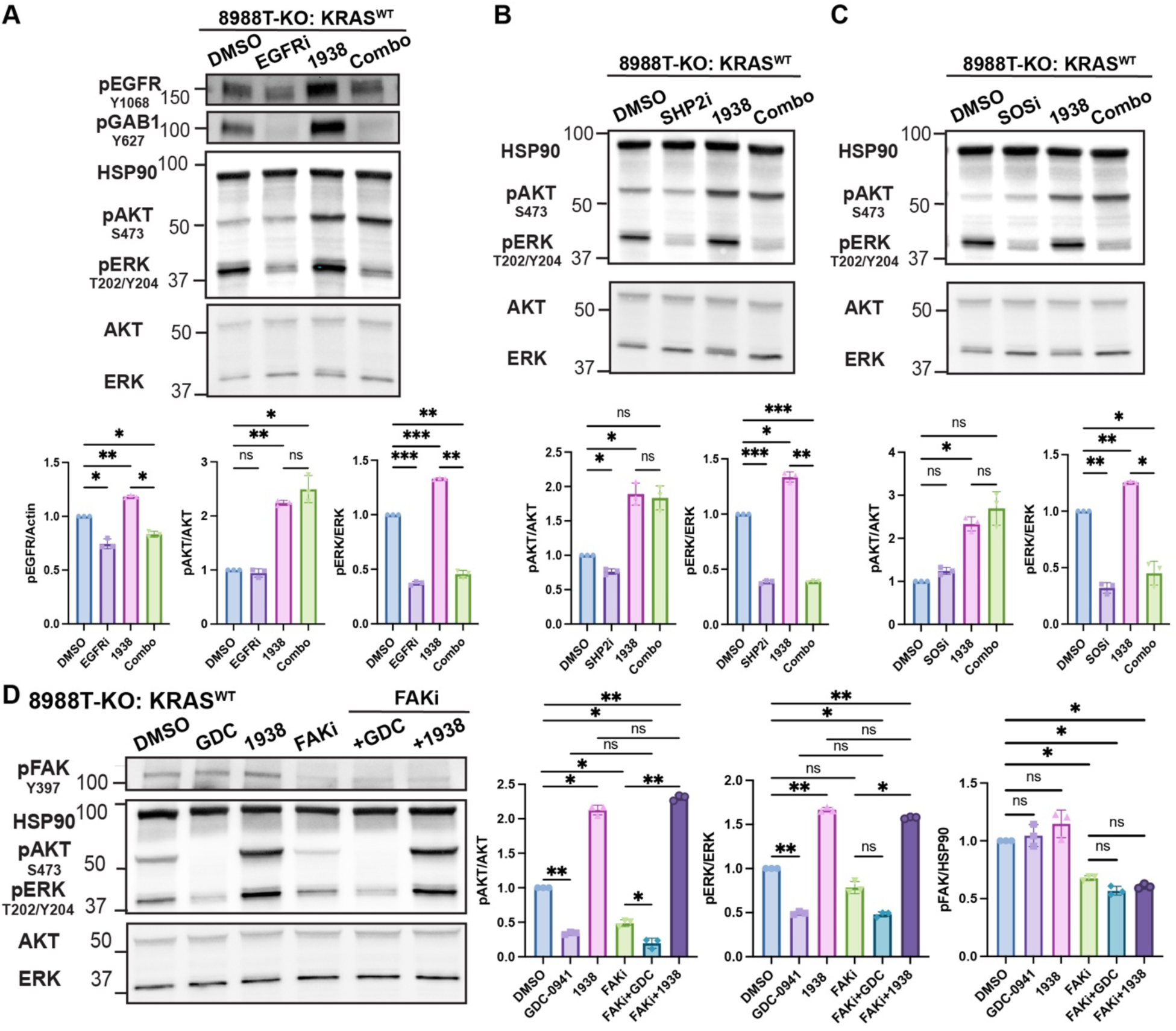
Molecular determinants of PI3K activity-dependent activation of RAS–MAPK signaling. (A) Representative immunoblots of phosphorylated EGFR (pEGFR^Y1068^), phosphorylated GAB1 (GAB1^Y627^), phosphorylated ERK1/2 (pERK^T202/Y204^), and phosphorylated AKT (pAKT^S473^) following treatment with DMSO (solvent), erlotinib (EGFRi, 5 μM), 1938 (1 μM), or their combination (Combo) for 0.5 hour in 8988T-KO: KRAS^WT^ cells. HSP90 is loading control. Bar graphs represent quantification (mean ± SD normalized to DMSO from n = 3 biological replicates). ns = non-significant, * p < 0.05, ** p < 0.01, *** p < 0.001, repeated measures ANOVA (rmANOVA) with Šidák’s post-hoc test. (B) Representative immunoblots of pERK1/2 and pAKT following treatment with DMSO (solvent), RMC-4550 (SHP2i, 2 μM), 1938 (1 μM), or their combination (Combo) for 0.5 hour in 8988T-KO: KRAS^WT^ cells. Bar graphs represent quantification (mean ± SD normalized to DMSO from n = 3 biological replicates). ns = non-significant, * p < 0.05, ** p < 0.01, *** p < 0.001, repeated measures ANOVA (rmANOVA) with Šidák’s post-hoc test. (C) Representative immunoblots of pERK1/2 and pAKT following treatment with DMSO (solvent), BI-3406 (SOSi, 1 μM), 1938 (1 μM), or their combination (Combo) for 0.5 hour in 8988T-KO: KRAS^WT^ cells. Bar graphs represent quantification (mean ± SD normalized to DMSO from n = 3 biological replicates). ns = non-significant, * p < 0.05, ** p < 0.01, *** p < 0.001, repeated measures ANOVA (rmANOVA) with Šidák’s post-hoc test. (D) Representative immunoblots of phosphorylated pERK1/2, pAKT, and phosphorylated FAK (pFAK^Y397^) following treatment with DMSO (solvent), defactinib (FAKi, 5 μM), 1938 (1 μM), or their combination (Combo) for 0.5 hour in 8988T-KO: KRAS^WT^ cells. Bar graphs represent quantification (mean ± SD normalized to DMSO from n = 3 biological replicates). ns = non-significant, * p < 0.05, ** p < 0.01, repeated measures ANOVA (rmANOVA) with Šidák’s post-hoc test.

### Genetic or pharmacologic PI3K activation drives resistance to KRAS inhibition

Recent studies suggest that PI3K activation may reduce the sensitivity of *RAS* mutant cancer cells to MAPK pathway inhibitors^17,54^. We hypothesized that PI3K activation could confer resistance to mutant KRAS inhibition in part by activating wild-type RAS signaling. To test this hypothesis, we treated AsPC1 and L3.3 PDAC cell lines – harboring KRAS^G12D^, the most common *KRAS* mutant allele in PDAC^55^ – with the KRAS^G12D^-specific inhibitor MRTX1133^56^ and observed that 1938 reduced sensitivity to MRTX1133 (**Figure 7A**). In contrast, 1938 had no effect on the sensitivity of these cell lines to the pan-RAS inhibitor RMC-6236, which is capable of inhibiting both mutant and wild-type RAS isoforms^57^ (**Figure 7B**). Importantly, PI3K inhibition (GDC-0941) markedly enhanced sensitivity to RAS inhibition across cell models and KRAS-targeted drugs, arguing that PI3K catalytic activity plays a role in mediating resistance to KRAS inhibition through both RAS-dependent and RAS-independent mechanisms (*e.g.*, AKT^7^).

**Figure 7.**
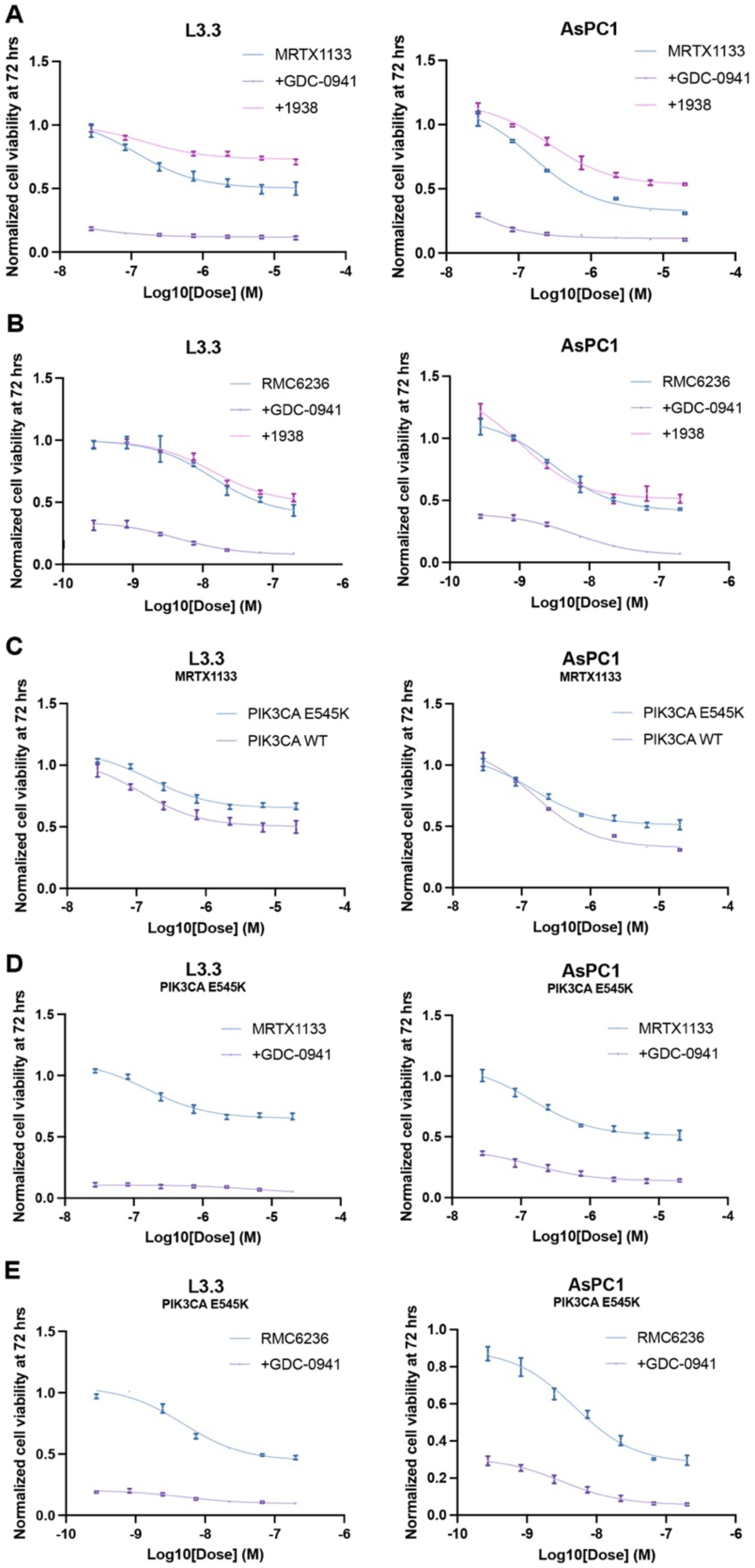
Genetic or pharmacologic PI3K activation drives resistance to KRAS inhibition. (A) Dose–response curves (three-parameter curve fit) of cell viability (normalized to DMSO control, mean ± SD, n = 3 technical replicates) of L3.3 and AsPC1 KRAS^G12D^ mutant PDAC cells treated with MRTX1133 +/- PI3K inhibitor (GDC-0941, 2 μM) or PI3K activator (1938, 1 μM). (B) Dose–response curves (three-parameter curve fit) of cell viability (normalized to DMSO control, mean ± SD, n = 3 technical replicates) of L3.3 and AsPC1 KRAS^G12D^ mutant PDAC cells treated with RMC-6236 +/- GDC-0941 (2 μM) or 1938 (1 μM). (C) Dose–response curves (three-parameter curve fit) of cell viability (normalized to DMSO control, mean ± SD, n = 3 technical replicates) of L3.3 and AsPC1 KRAS^G12D^ mutant PDAC cells transduced with PIK3CA^WT^ or PIK3CA^E545K^ and treated with MRTX1133. (D) Dose–response curves (three-parameter curve fit) of cell viability (normalized to DMSO control, mean ± SD, of n = 3 technical replicates) of L3.3 and AsPC1 KRAS^G12D^ mutant PDAC cells transduced with PIK3CA^E545K^ and treated with MRTX1133 +/- GDC-0941 (2 μM). (E) Dose–response curves (three-parameter curve fit) of cell viability (normalized to DMSO control, mean ± SD, n = 3 technical replicates) of L3.3 and AsPC1 KRAS^G12D^ mutant PDAC cells transduced with PIK3CA^E545K^ and treated with RMC-6236 +/- GDC-0941 (2 μM).

To further elucidate the clinical relevance of PI3K-driven resistance, we examined data from the CodeBreaK100 trial, which identified PIK3CA-activating mutations E542K and E545K – both mapping to the helical domain of p110α – as potential contributors to resistance to the KRAS^G12C^ inhibitor sotorasib in PDAC patients^12^. Strikingly, lentivirus-mediated expression of either PIK3CA^E542K^ or PIK3CA^E545K^ was sufficient to increase both pAKT and pERK levels in 8988T-KO cells, phenocopying the effects of 1938-mediated PI3K activation (**Figure S6A**). To determine whether these mutations could similarly promote resistance to KRAS inhibition, we expressed PIK3CA^E545K^ in AsPC1 and L3.3 cells and assessed their response to KRAS^G12D^ inhibition. Consistently, PIK3CA^E545K^ reduced the sensitivity of both cell lines to MRTX1133 (**Figure 7C**). Conversely, PI3K inhibition sensitized these cells to MRTX1133 (**Figures 7D and 7E**) and led to a stronger suppression of pERK (**Figures S6B and S6C**). Collectively, our findings highlight PI3K activation as a key driver of resistance to allele-specific KRAS inhibition, mediated, in part, by the activation of wild-type RAS–MAPK signaling. These results underscore the potential of combining PI3K and KRAS inhibitors as an effective therapeutic strategy to overcome resistance in *KRAS* mutant PDAC.

## DISCUSSION

The clinical success of KRAS-targeted therapies has been tempered by the rapid onset of resistance. While secondary genetic alterations have been implicated in approximately 50% of cases resistant to KRAS^G12C^ inhibitors, it is increasingly evident that non-genetic mechanisms – which are often more rapid and plastic – play an important role in driving resistance to KRAS-targeted therapies^58^. Addressing these adaptive, non-genetic processes is critical for developing more durable therapeutic strategies. Here, we identify a novel mode of non-genetic resistance to KRAS inhibition in PDAC cells, in which PI3K sustains wild-type RAS–MAPK signaling by orchestrating GAB1-mediated signaling complex formation at the plasma membrane (**Figure S7**). This challenges the conventional linear model of RAS signaling and suggests that the spatial organization of signaling platforms can sustain oncogenic output even when mutant KRAS is inhibited.

Although PI3K has classically been viewed as a downstream effector of RAS^59,60^, several studies have reported its capacity to activate MAPK signaling^22,24,50,61^. For example, isoform-specific and pan-PI3K inhibitors transiently reduced RAS–MAPK activation in RAS^WT^ breast cancer cells harboring *HER2* amplifications or *PIK3CA* mutations^22^. These findings align closely with our data, which show a strong correlation between pERK and pAKT suppression in RAS^WT^ – but not mutant – models (**Figures 1 and 2**). Importantly, our studies overcome many limitations of prior observational and mechanistic work which used non-specific PI3K inhibitors (*e.g.,* wortmannin^62^, LY294002^63^), relied on exogenous ligand activation of RTKs, lacked spatial considerations in mechanistic analyses, and contained insufficient functional analyses linking PI3K to RAS activation. Furthermore, our data – leveraging recently developed isoform-specific inhibitors – argue for functional redundancy between class IA PI3K isoforms in MAPK regulation, as has been observed in PI3K-mediated resistance to BRAF/MEK inhibitors in melanoma^64^. Finally, our findings suggest that PI3K can be either an upstream regulator or downstream effector of RAS depending on *RAS* mutational status (**Figure S7**).

Our results underscore how PI3K signals to wild-type RAS to activate MAPK signaling. PI3K serves as a master spatial organizer of oncogenic signaling through modulation of membrane-associated lipids and scaffold protein networks including GAB1 (**Figures 3-5**). This signalosome assembles primarily outside focal adhesions, suggesting that PI3K-driven RAS activation requires co-recruitment of additional positive regulators (EGFR, SHP2, GRB2, SOS1) within a distinct spatial niche. Of note, EGFR phosphorylates GAB1 at Y627, even in the absence of exogenous ligand, suggesting that basal activation of EGFR is sufficient for basal RAS–MAPK activation by PI3K or in the context of allosteric PI3K activation. These data are consistent with prior studies in which wild-type RAS compensates for oncogenic KRAS inhibition, particularly through feedback activation involving RTKs (including EGFR)^15,65^, SHP2^66,67^, and SOS1^68^. Our results argue that PI3K drives the assembly of these EGFR/SHP2/GRB2/SOS1 signaling complexes to induce resistance to mutant KRAS inhibition (**Figures 4E-H, 6, and 7**), though the structural details of these multiprotein complexes remain to be elucidated. Importantly, this PI3K-dependent bypass mechanism has direct clinical relevance. Activating mutations in *PIK3CA* have been observed in PDAC patients who relapse following treatment with KRAS^G12C^ inhibitors^12^. Nonetheless, PI3K inhibition enhances sensitivity to KRAS inhibition across both *PI3KCA* mutant and wild-type cell models (**Figure 7**), supporting the rational development of combinatorial therapeutic strategies involving PI3K and KRAS inhibitors for PDAC.

### Limitations of the Study

Most experiments defining the mechanistic basis for PI3K-mediated RAS–MAPK signaling were performed in cell culture models, which do not fully capture the spatial and biochemical complexity of native tissue environments. Although validation using *in vivo* models would strengthen the physiologic relevance of our conclusions, such studies remain technically challenging due to the transient nature of MAPK pathway suppression following PI3K inhibition, variability in pharmacokinetics and specificity of PI3K inhibitors, and the confounding effects of insulin feedback^69^. We focused our study on PDAC due to the high frequency of *KRAS* mutations^1^, the limited effective therapeutic options making KRAS inhibition more attractive, and the availability of *KRAS*-less cells we have generated enabling the study of KRAS-related signaling mechanisms in a clean background^7,28^. Whether our findings apply to other cancers is not known, especially as response to PI3K inhibitors can vary by cancer type^70^. However, previous studies have shown synergistic anti-tumor activity between KRAS and PI3K inhibitors in preclinical lung cancer models^71^, suggesting similar mechanisms may still apply. Finally, while GAB1 emerged as a key PI3K-dependent scaffold, our proteomic analyses identified additional adaptors and docking proteins – many with known PH domains – that may contribute to RAS–MAPK signaling (**Figure 5A**). The functional roles of these proteins, including potential redundancies or cooperative interactions, remain to be fully explored.

## RESOURCE AVAILABILITY

### Lead contact

Further information and requests for resources and reagents should be directed to and will be fulfilled by the lead contact, Mandar Deepak Muzumdar.

### Materials availability

Plasmids generated in this study will be deposited in Addgene. All unique/stable reagents generated in this study are available from the lead contact with a completed materials transfer agreement.

### Data and code availability

All raw mass spectrometry data were deposited in the ProteomeXchange Consortium via the Proteomics Identifications Database (PRIDE)^72,73^ and will be publicly available as of the date of publication. This paper does not report original code. Any additional information required to reanalyze the data reported in this paper is available from the lead contact upon request.

## ACKNOWLEDGEMENTS

We thank the Muzumdar lab members for helpful discussions and feedback; Drs. Mark Lemmon and David Stern for critical reading of the manuscript; the Yale West Campus Imaging Core, Yale West Campus Analytic Core, the Yale Keck DNA Sequencing Core, MGH CCIB DNA Core, and MIT BioMicroCenter for microscopy, proteomics, and DNA sequencing; Drs. Gordon Mills, Tyler Jacks, Dafna Bar-Sagi, Alice Ting, Jared Toettcher, the National Cancer Institute (NCI) RAS Initiative, and Dr. Clotilde Calderwood and the Yale Cancer Center (YCC) Functional Genomics Core for plasmid constructs. C.S.M. was supported by a National Science Foundation (NSF) Graduate Research Fellowship. C.F.R. was supported by a postdoctoral fellowship through the Yale Cancer Biology Training Program (T32-CA193200) and is supported by an NCI Research Supplement (R01-CA276108-02S2). M.B. acknowledges support from the National Institute of General Medical Sciences (NIGMS: R35GM147095). Y.L. acknowledges support from the NIGMS (R01GM137031, RM1GM149406). M.D.M. acknowledges support from an NIH Director’s New Innovator Award (DP2-CA248136), NCI Mentored Clinical Scientist Research Career Development Award (K08-CA2080016), American Cancer Society Institutional Research Grant (IRG 17-172-57), Lustgarten Foundation Therapeutics-Focused Research Program, NCI R01-CA276108, and in part, the Yale Comprehensive Cancer Center Support Grant (P30-CA016359). The content is solely the responsibility of the authors and does not necessarily represent the official views of the National Institutes of Health.

## AUTHOR CONTRIBUTIONS

X.G.: conceptualization, data curation, formal analysis, investigation, methodology, visualization, writing-original draft; J.S.: data curation, formal analysis, investigation, visualization, writing-review & editing; W.L.: formal analysis, investigation, methodology, writing-review & editing; C.S.M.: data curation, formal analysis, visualization, writing-review & editing; C.F.R.: formal analysis, visualization, writing-review & editing; M.B.: methodology, supervision, writing-review & editing; Y.L.: methodology, resources, supervision, writing-review & editing; M.D.M.: conceptualization, data curation, formal analysis, funding acquisition, investigation, project administration, resources, supervision, visualization, writing-original draft.

## SUPPLEMENTAL INFORMATION

**Table S1. Protein proximity labeling proteomics data, related to Figures 4 and 5**

**Table S2. Differential biotinylated protein abundance with PI3K activity modulation, related to Figures 4 and 5**

**Table S3. Brunello CRISPR screen MaGECK ranks, related to Figure 4**

## SUPPLEMENTAL FIGURE LEGENDS

**Figure S1.**
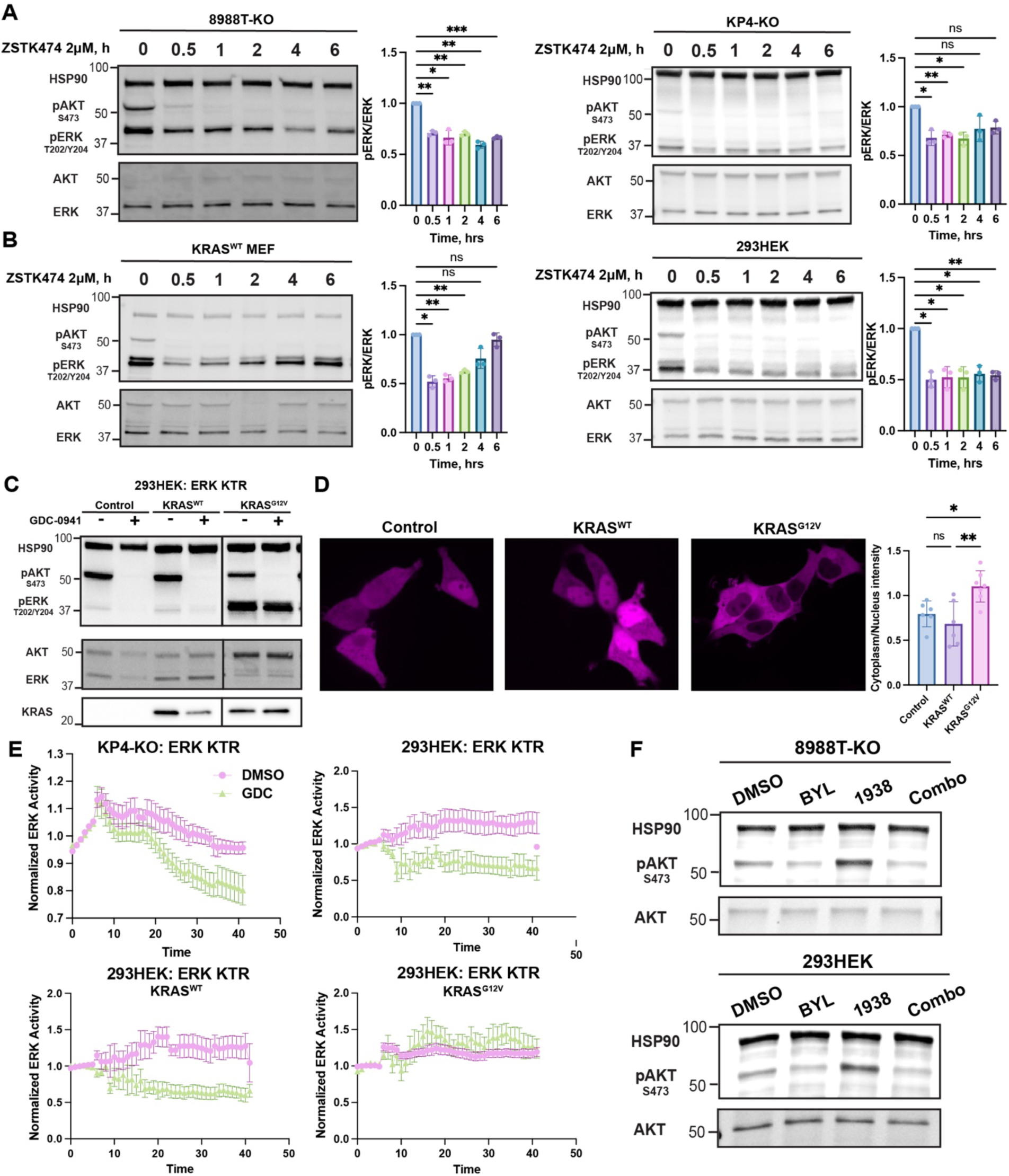
PI3K regulates MAPK signaling in wild-type RAS cells independent of p110 isoforms, related to Figure 1. (A) Representative immunoblots showing the effect of PI3K inhibition (ZSTK474, 2 μM) over a 6-hour time course on phosphorylated ERK1/2 (pERK^T202/Y204^) and phosphorylated AKT (pAKT^S473^) in KRAS deficient PDAC clones (8988T-KO, KP4-KO). HSP90 serves as loading controls. Bar graphs quantify pERK/ERK and pAKT/AKT levels normalized to time 0 (mean ± SD, n = 3 biological replicates). ns = not significant, * p < 0.05, ** p < 0.01 *** p < 0.001, repeated measures ANOVA (rmANOVA) with Dunnett’s post-hoc test. (B) Representative immunoblots showing the effect of PI3K inhibition (ZSTK474, 2 μM) over a 6-hour time course on pERK1/2 and pAKT in non-malignant cells (293HEK, KRAS^WT^ MEF cells). Bar graphs quantify pERK/ERK and pAKT/AKT levels normalized to time 0 (mean ± SD, n = 3 biological replicates). ns = not significant, * p < 0.05, ** p < 0.01, rmANOVA with Dunnett’s post-hoc test. (C) Immunoblots showing the effects of PI3K inhibition (GDC-0941, 2 μM) on pERK1/2 and pAKT in 293HEK cells transfected with real-time ERK kinase translocation reporter (ERK-KTR) and KRAS^WT^ or KRAS^G12V^ constructs. pERK is robustly increased with KRAS^G12V^, and is resistant to PI3K inhibition, unlike KRAS^WT^ and control (not transfected with KRAS construct, thus representing endogenous signaling). HSP90 is loading control. (D) Representative images of ERK-KTR reporter in 293HEK: ERK-KTR cells in (**C**) showing cytoplasmic redistribution of ERK-KTR in KRAS^G12V^-expressing cells indicating elevated ERK activity. Bar graphs show quantification of ERK activity (cytoplasmic-to-nuclear fluorescence intensity ratio, mean ± SD, n = 6-7 biological replicates per group). ns = not significant, * p < 0.05, ** p < 0.01, one-way ANOVA with Tukey’s post-hoc test. (E) Quantification of ERK activity (as in (**D**)) normalized to average baseline intensity by time-lapse imaging of KP4-KO and 293HEK cells (untransfected or transfected with KRAS^WT^ or KRAS^G12V^) following treatment with GDC-0941 vs. DMSO solvent showed a rapid decrease in ERK activity in RAS wild-type cells (n=5-10 cells per group).

**Figure S2.**
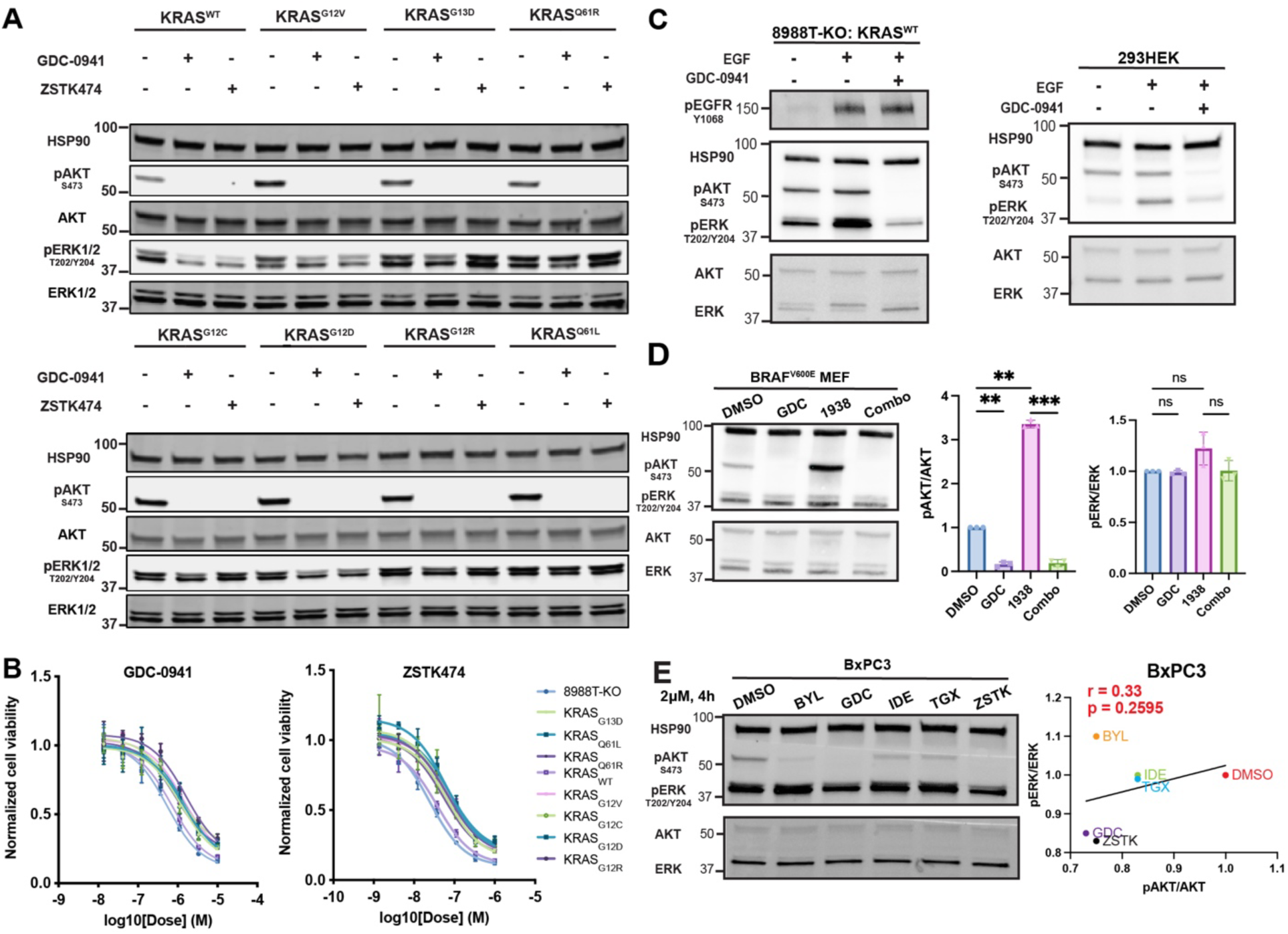
PI3K selectively regulates wild-type RAS activity but not mutant RAS, related to Figure 2. (A) Representative immunoblots of phosphorylated ERK1/2 (pERK^T202/Y204^) and phosphorylated AKT (pAKT^S473^) in 8988T-KO cells reconstituted with various KRAS mutants (G12V, G13D, Q61R, G12C, G12D, G12R, Q61L) or KRAS^WT^ treated with GDC-0941 or ZSTK474 (2 μM) for 4 hours. HSP90 is loading control. (B) Dose–response curves (three-parameter curve fit) of cell viability (normalized to DMSO control, mean ± SD, n = 3 technical replicates) for GDC-0941 and ZSTK474 of cell lines in **(A)**, demonstrating that mutant KRAS (but not KRAS^WT^) decreased PI3K inhibitor sensitivity irrespective of mutant variant. (C) Representative immunoblots of pERK1/2 and pAKT in 8988T-KO KRAS^WT^ cells and HEK293 cells stimulated with EGF (100 ng/mL) in the presence or absence of the PI3K inhibitor GDC-0941 (2 μM) for 30 minutes. (D) Representative immunoblots and quantification of pERK1/2 and pAKT following treatment with DMSO (solvent), 2 μM GDC-0941 (GDC), 1 μM PI3K activator 1938, or their combination for 0.5 hour in BRAF^V600E^ MEFs. Bar graphs represent mean ± SD normalized to DMSO from n = 3 biological replicates. ns = non-significant, ** p < 0.01, *** p < 0.001, repeated measures ANOVA (rmANOVA) with Šidák’s post-hoc test. (E) Immunoblots showing phosphorylation levels of pERK1/2 and pAKT in BxPC3 PDAC cells (KRAS^WT^; BRAF^V600E^) following 4-hour treatment with isoform-selective PI3K inhibitors (2 μM each): BYL-719 (BYL, p110α), GDC-0941 (GDC, pan-class IA), idelalisib (IDE, p110δ), TGX-221 (TGX, p110β), and ZSTK474 (ZSTK, pan-class IA). Correlation between pAKT/AKT and pERK/ERK across inhibitor treatments was assessed, indicating that PI3K signaling activity (pAKT) is not associated with MAPK output (pERK). Pearson correlation coefficient (r) and associated p-value are shown,

**Figure S3.**
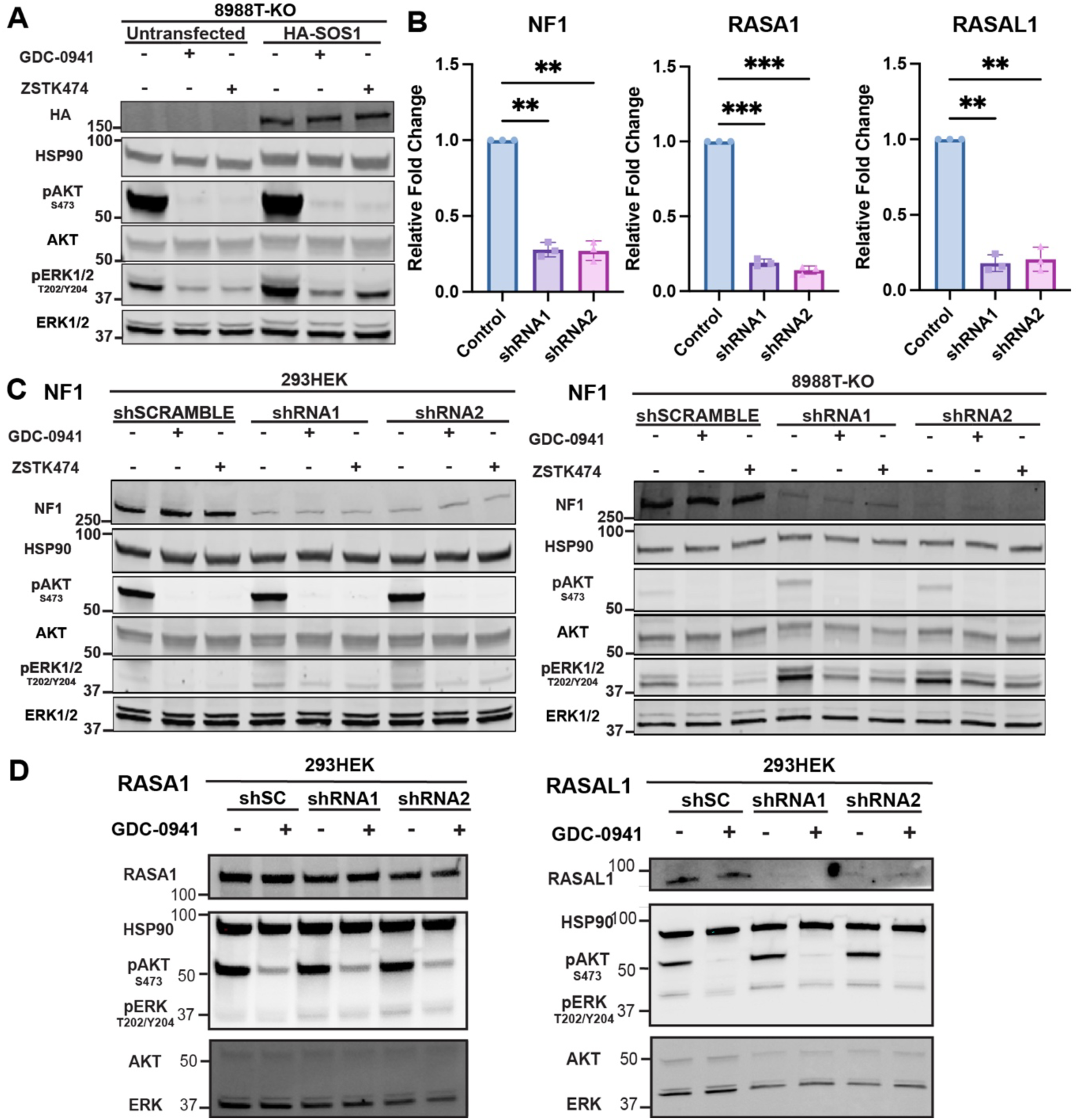
PI3K-mediated wild-type RAS signaling is independent of canonical GEF and GAP expression, related to Figure 3. (A) Immunoblots of phosphorylated ERK1/2 (pERK^T202/Y204^) and phosphorylated AKT (pAKT^S473^) in 8988T-KO cells transduced with HA-tagged SOS1 and treated with GDC-0941 (2 μM) or ZSTK474 (2 μM) for 4 hours. HSP90 is loading control. (B) Quantification of knockdown of GAPs NF1, RASA1, and RASAL1 by quantitative RT-PCR. Bar graph shows relative mRNA levels normalized to control (mean ± SD, n = 3 biologic replicates). ** p < 0.01, *** p < 0.001, repeated measures ANOVA (rmANOVA) with Dunnett’s test. (C) Immunoblots of pERK1/2 and pAKT in 293HEK and 8988T-KO cells subject to stable knockdown of NF1 using two independent shRNAs and treated with GDC-0941 or ZSTK474 (2 μM, 4 hours). (D) Immunoblots of pERK1/2 and pAKT in 293HEK cells subject to stable knockdown of RASA1 or RASAL1 using two independent shRNAs and treated with GDC-0941 (2 μM, 4 hours).

**Figure S4.**
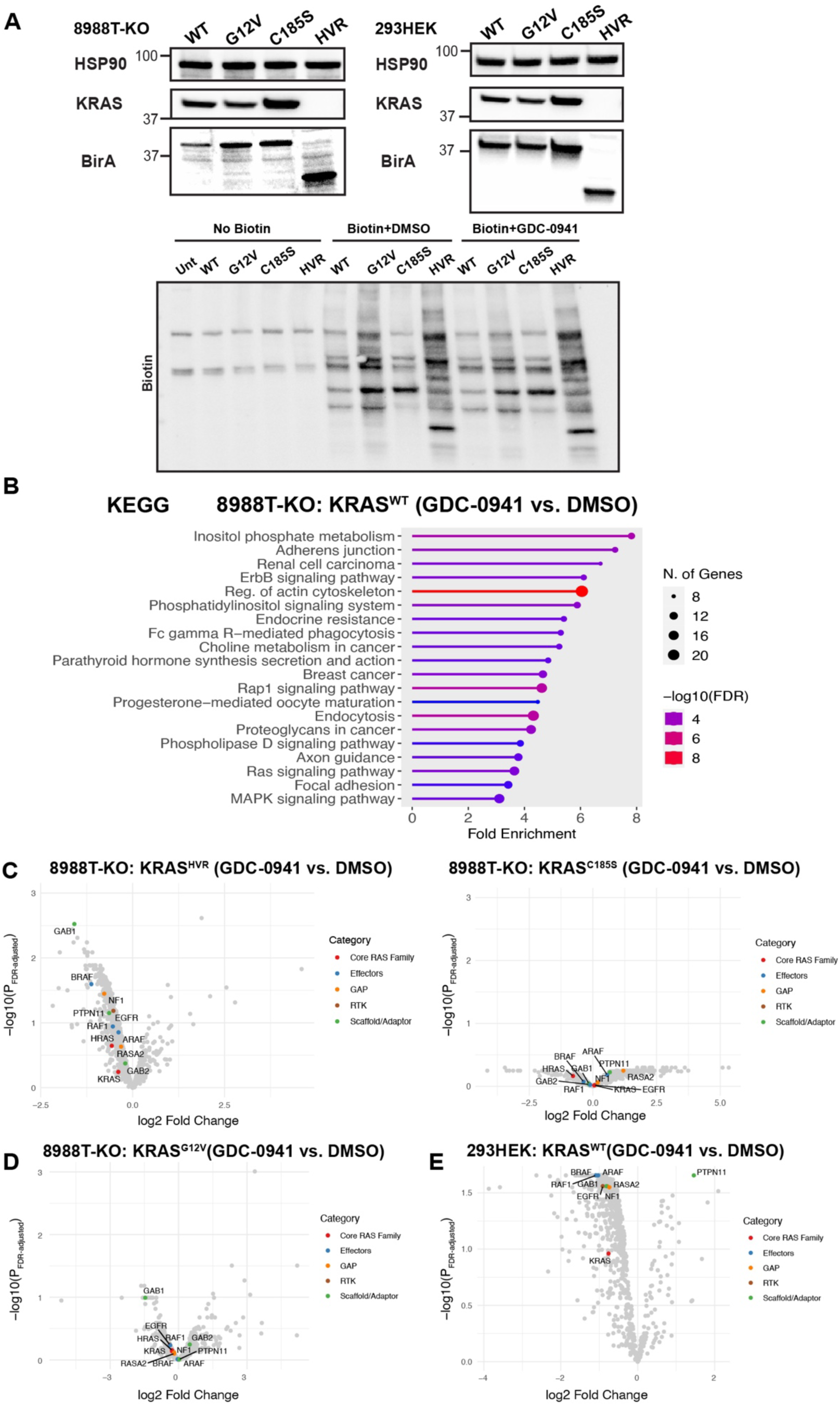
Protein proximity labeling identifies PI3K-dependent KRAS interactors, related to Figure 4. (A) Immunoblots showing proximity-based biotinylation profiles from PDAC and 293HEK cells expressing BirA-tagged KRAS variants (WT, G12V, C185S, and HVR) following treatment with DMSO or GDC-0941 (2 μM). Biotin vs. no biotin confirms enrichment of specific proteins in cells harboring BirA constructs. (B) KEGG pathway enrichment analysis of decreased biotinylated proteins in 8988T-KO: KRAS^WT^ cells treated with GDC-0941 (vs. DMSO control) reveals significant enrichment in RAS-MAPK, PI3K-AKT, focal adhesion, and actin cytoskeleton pathways. (C) Volcano plots (log2 fold-change (LFC) vs. −log10(adjusted p-value)) showing differential biotinylated protein enrichment between GDC-0941 and DMSO conditions 8988T-KO: KRAS^HVR^ and 8988T-KO: KRAS^C185S^ PDAC cells. (D) Volcano plot (LFC vs. −log10(adjusted p-value)) showing differential biotinylated protein enrichment between GDC-0941 and DMSO conditions 8988T-KO: KRAS^G12V^ cells. (E) Volcano plot (LFC vs. −log10(adjusted p-value)) showing differential biotinylated protein enrichment between GDC-0941 and DMSO conditions 293HEK: KRAS^WT^ cells, showing similar results in decreased proteins to 8988T-KO: KRAS^WT^ cells.

**Figure S5.**
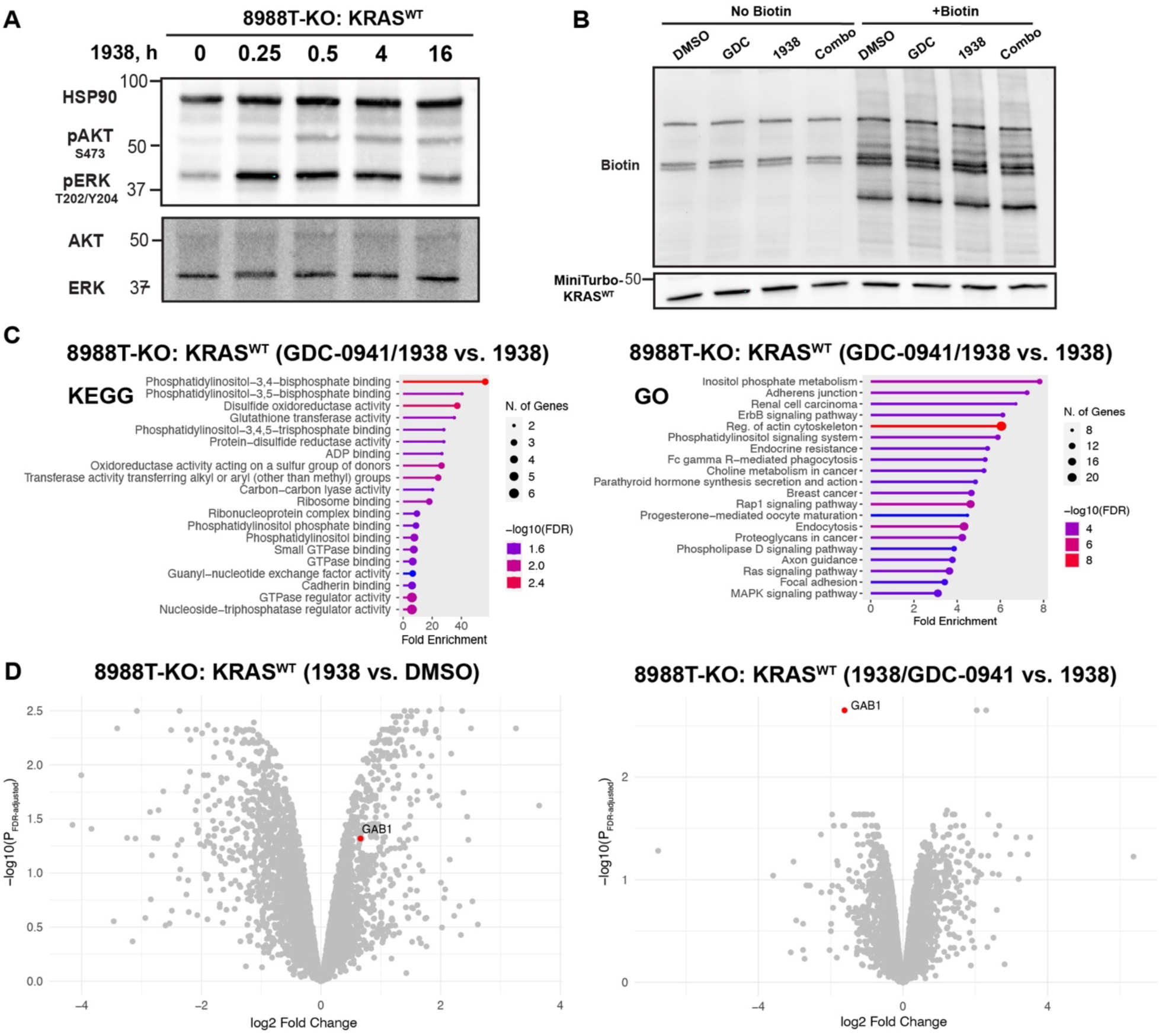
Protein proximity labeling using MiniTurbo identifies PI3K-dependent KRAS interactors, related to Figure 5. (A) Immunoblots showing the transient effects of PI3K activation (1938, 1 μM) over a 16-hour time course on phosphorylated ERK1/2 (pERK^T202/Y204^) and phosphorylated AKT (pAKT^S473^) in 8988T-KO cells. (B) Immunoblots showing proximity-based biotinylation profiles from 8988T-KO: MiniTurbo-KRAS^WT^ cells following treated with DMSO, GDC-0941 (2 μM), 1938 (1 μM), or both. Biotin vs. no biotin confirms enrichment of specific proteins in cells harboring MiniTurbo-KRAS. (C) KEGG and Gene Ontology (GO) pathway enrichment analysis of decreased biotinylated proteins in 8988T-KO: MiniTurbo-KRAS^WT^ cells co-treated with GDC-0941 and 1938 (vs. 1938 alone) reveals significant enrichment in phosphoinositide binding and metabolism, GTPase regulation, RAS–MAPK signaling, and focal adhesion proteins. (D) Volcano plots (log2 fold-change (FC) vs. −log10(adjusted p-value)) of 8988T-KO: MiniTurbo-KRAS^WT^ cells showing that GAB1 is significantly enriched upon 1938 treatment and is the most significantly reduced protein with concurrent GDC-0941 treatment.

**Figure S6.**
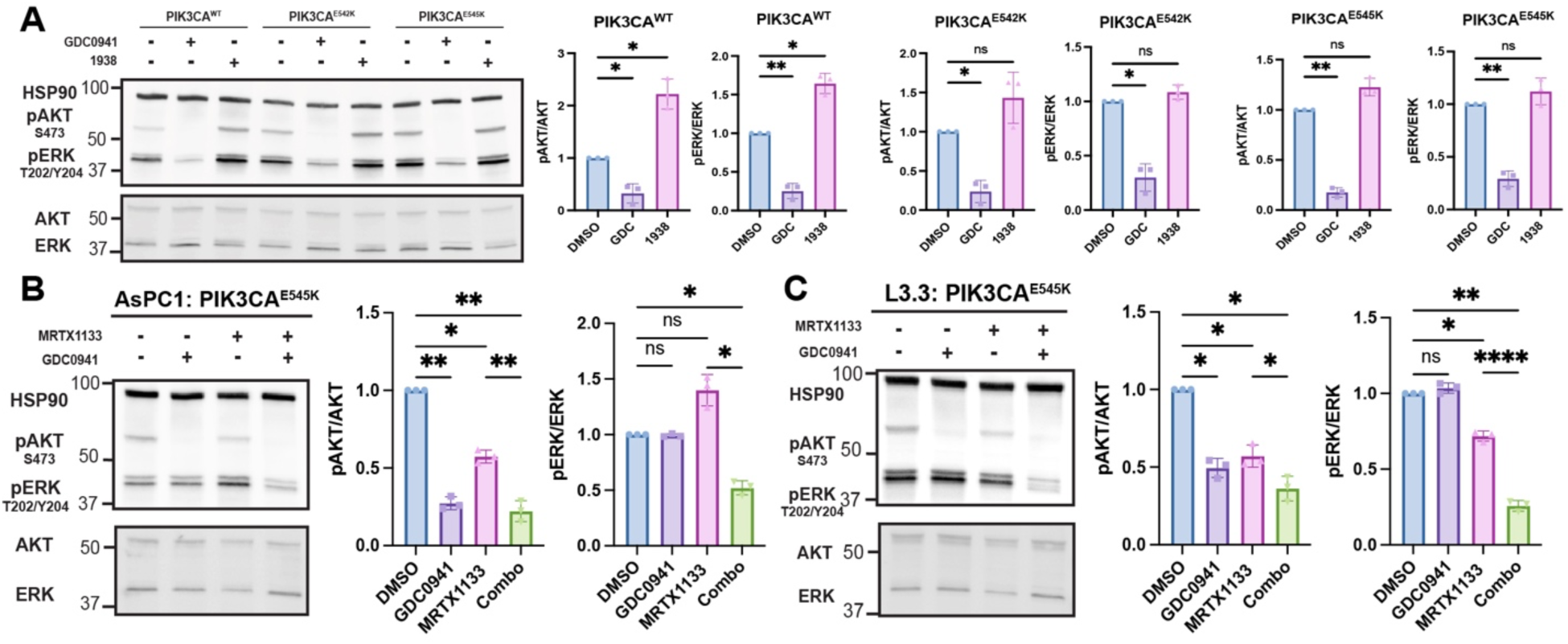
Genetic or pharmacologic PI3K activation drives resistance to KRAS inhibition, related to Figure 7. (A) Representative immunoblots of phosphorylated ERK1/2 (pERK^T202/Y204^) and phosphorylated AKT (pAKT^S473^) in 8988T-KO cells expressing PIK3CA^WT^ or mutant PIK3CA (E542K or E545K) following treatment with DMSO, GDC-0941 (2 μM), or 1938(1 μM). HSP90 is loading control. Bar graphs represent quantification of pAKT/AKT and pERK/ERK normalized to DMSO (mean ± SD, n = 3 biological replicates). ns = non-significant, * p < 0.05, ** p < 0.01, repeated measures ANOVA (rmANOVA) with Dunnett’s test. (B) Representative immunoblots of pERK1/2 and pAKT in AsPC1: PIK3CA^E545K^ cells treated with MRTX1133 (100 nM), GDC-0941 (2 μM), or both. Bar graphs represent quantification of pAKT/AKT and pERK/ERK normalized to DMSO (mean ± SD, n = 3 biological replicates). ns = non-significant, * p < 0.05, ** p < 0.01, rmANOVA with Šidák’s post-hoc test). (C) Representative immunoblots of pERK1/2 and pAKT in L3.3: PIK3CA^E545K^ cells treated with MRTX1133 (100 nM), GDC-0941 (2 μM), or both. Bar graphs represent quantification of pAKT/AKT and pERK/ERK (mean ± SD normalized to DMSO for each construct, n = 3 biological replicates). * p < 0.05, ** p < 0.01, **** p < 0.0001, rmANOVA with Šidák’s post-hoc test.

**Figure S7.**
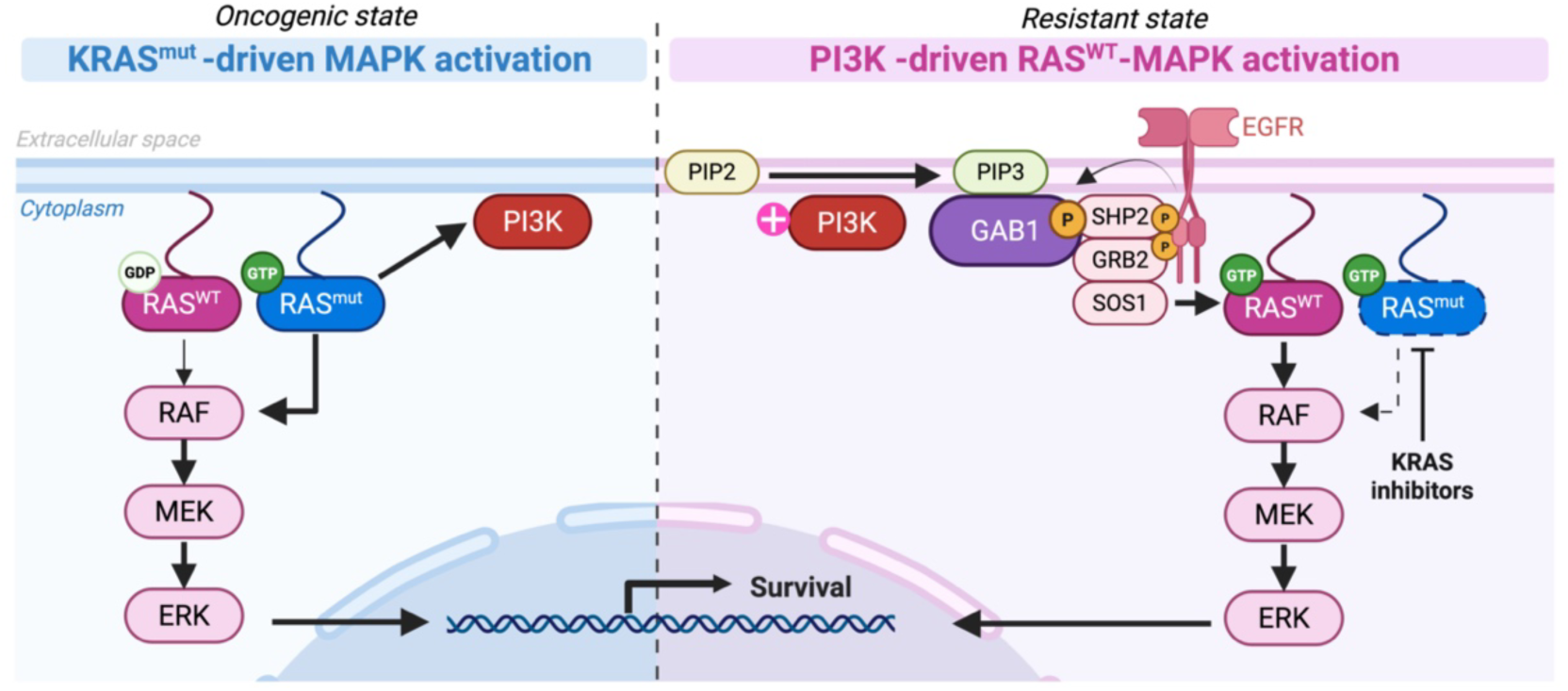
Model of PI3K-mediated wild-type RAS–MAPK signaling as a bypass mechanism to KRAS inhibition, related to Figures 4-7. In the oncogenic state, mutant KRAS drives both MAPK and PI3K signaling. Upon KRAS inhibition, PI3K facilitates the assembly of GAB1/GRB2/EGFR/SHP2 signaling complexes, which sustain wild-type RAS–MAPK signaling to promote cell survival and thereby conferring resistance to mutant KRAS-targeted therapy. Created with BioRender.com.

## METHODS

### EXPERIMENTAL MODEL AND STUDY PARTICIPANT DETAILS

#### Cell lines and cell culture conditions

Established human PDAC cell lines (8988T, BxPC3, KP-4, AsPC1, and L3.3) and 293T cells (293HEK) were sourced from DSMZ-Germany, American Type Culture Collection (ATCC), and RIKEN. *Ras*-less mouse embryonic fibroblasts (MEFs) reconstituted with *RAS* and *BRAF* variants were acquired from the National Cancer Institute (NCI) RAS initiative. *KRAS*-deficient cell lines (8988T-KO, KP4-KO, and 8988T clones used in CRISPR screens) were generated by CRISPR knockout of *KRAS*, as previously described^7^. Clones were confirmed to retain (intact) or lack (deficient) KRAS protein expression by immunoblotting and harbor no mutation (intact) or out-of-frame mutations (deficient) by Sanger sequencing or amplicon sequencing by the MGH CCIB DNA Core. Isogenic cell lines were generated by reconstituting *KRAS* variants in 8988T-KO cells through lentiviral transduction. All cell lines were confirmed negative for mycoplasma using PCR testing. All cell lines except for AsPC1 and L3.3 were maintained in DMEM (Corning Cellgro) supplemented with 10% fetal bovine serum (FBS) (Thermo Fisher Scientific) and 1% penicillin/streptomycin (Thermo Fisher Scientific). AsPC1 and L3.3 cells were cultured in RPMI-1640 medium (Thermo Fisher Scientific) supplemented with 10% FBS, 1% penicillin-streptomycin, 25 mM glucose (Thermo Fisher Scientific), 1 mM sodium pyruvate (Thermo Fisher Scientific), and 10 mM HEPES (Thermo Fisher Scientific). For all treatment studies, cells were plated in fresh media and then drugs were added to the media the next day. Treatment used in the study include: GDC-0941 (Selleckchem, S1065, 2 μM), ZSTK474 (Selleckchem, S1072, 2 μM), defactinib (Selleckchem, S7654, 5 μM), TGX-221 (Selleckchem, S1169, 2 μM), idelalisib (Selleckchem, S2226, 2 μM), BYL-719 (Selleckchem, S2814, 2 μM), RMC-6236 (Selleckchem, E1597), MRTX1133 (MedChemExpress, HY-134813), erlotinib (Selleckchem, S7786, 5 μM), RMC-4550 (Selleckchem, S8718, 2 μM), BI-3406 (Selleckchem, S8916, 1 μM), MK-2206 (Selleckchem, S1078, 2 μM), UCL-TRO-1938 (Selleckchem, E1579, 1 μM), and EGF (R&D Systems, 236-EG, 10 ng/ml) were used.

#### Plasmid construction

Lentiviral plasmids were assembled using Gibson Assembly (NEB, E2611), as previously described^74^. Briefly, lentiviral backbone vector (LV 1-5, Addgene #68411) was linearized by restriction enzyme digestion at designated sites. Inserts were PCR-amplified with Q5 Hot Start High-Fidelity DNA Polymerase (NEB, M0493) using primers with 20–30 bp overlaps. PCR products were gel-purified (Qiagen, 28704) and assembled at 50°C for 1 hour. The assembly mixtures were transformed into Stbl3 competent E. coli (Invitrogen, C737303) and selected on antibiotic-containing agar plates. Individual positive clones were cultured overnight in Terrific Broth (Thermo Fisher Scientific). Plasmids were purified using a miniprep kit (Qiagen, 27106) and validated by Sanger sequencing performed by the Keck Biotechnology Resource Laboratory at Yale. Plasmids used for PCR amplification of wild-type/mutant RAS isoforms were obtained from the NCI RAS Initiative. Plasmids for cDNA amplification of fluorescent proteins (*GFP* and *mCherry*), *BirA*, *MiniTurbo*, *SOS1*, and *PIK3CA* (WT/E542K/E545K) were obtained from Addgene: pET21a-BirA (#20857), miniTurbo-His6_pET21a (#107178), HA-SOS1 (#32920), pHAGE-PIK3CA (#116771), pHAGE-PIK3CA-E542K (#116479), pHAGE-PIK3CA-E545K (#116485). GAB1 ORF was obtained through the Yale Cancer Center (YCC) Functional Genomics Core and sourced from the DNASU Plasmid Repository.

#### Lentiviral production and transduction

To produce lentivirus, we transfected plasmids that contained lentiviral backbone, packaging vector (psPAX2, Addgene #12260), and envelope vector (VSV-G, Addgene #8454) into 293HEK cells using TransIT-LT1 (Mirus Bio). Supernatants were collected at 48- and 72-hours post-transfection and applied to target cells supplemented with 8 μg/mL polybrene (EMD Millipore). To isolate cell populations expressing fluorescently tagged proteins, cells were sorted based on fluorophore intensity using a FACSMelody cell sorter (BD Biosciences). Expression of the fusion proteins was confirmed by immunoblotting.

#### RNA interference-mediated gene knockdown

Short hairpin RNA (shRNA) plasmids targeting specific genes, along with a non-targeting scramble control, were obtained through the YCC Functional Genomics Core and sourced from the MISSION lentiviral shRNA library (Sigma). shRNA sequences included:

NF1-shRNA1: 5’-CCATGTTGTAATGCTGCACTT-3’
NF1-shRNA2: 5’-GCCAACCTTAACCTTTCTAAT-3’
RASA1-shRNA1: 5’-GCTGCCTAACTTATCCATCTT-3’
RASA1-shRNA2: 5’- CCTGGCGATTATTCACTTTAT-3’
RASAL1-shRNA1: 5’- GCCATTGTTCGAGAAGGCTAT-3’
RASAL1-shRNA2: 5’- CCTGTGATTAGCCGTGTCTTT-3’
PTEN-shRNA: 5’-AGGCGCTATGTGTATTATTAT-3’
Scramble-shRNA: 5’-CAACAAGATGAAGAGCACCAA-3’

293HEK cells were transfected with individual shRNA plasmids using TransIT-LT1 Transfection Reagent (Mirus Bio) according to the manufacturer’s instructions. 8988T-KO cells were transduced with shRNA lentivirus and selected with puromycin (2 μg/mL) for three days. At 72 hours post-transfection or post-transduction, cells were treated with either DMSO (vehicle control) or 2 μM GDC-0941 for 30 minutes. Cells were harvested and lysed for immunoblotting and/or quantitative real-time PCR to assess knockdown efficiency and downstream signaling effects.

#### Quantitative Real-Time PCR (qRT-PCR)

Total RNA was extracted using the RNeasy Mini Kit (Qiagen) following the manufacturer’s protocol, including on-column DNase I treatment to eliminate genomic DNA contamination. RNA concentration and purity were assessed using a NanoDrop spectrophotometer (Thermo Fisher Scientific). cDNA synthesis was performed using the High-Capacity cDNA Reverse Transcription Kit (Thermo Fisher Scientific, 4368814) with 1 μg of total RNA per reaction, according to the manufacturer’s instructions. Quantitative PCR was performed using TaqMan™ Universal PCR Master Mix (Thermo Fisher, Scientific, 4304437) and gene-specific FAM probes (Thermo Fisher Scientific) on a CFX Opus 384 Real-Time PCR System (Bio-Rad) with the following probes: GAPDH(Hs99999905_m1), RASA1(Hs00243115_m1), RASAL1(Hs00183013_m1), NF1(Hs01035108_m1). Gene expression was normalized to GAPDH as the internal control.

#### Immunoblotting

Whole cell lysates were prepared in Pierce® RIPA buffer (Thermo Fisher Scientific) supplemented with 1:100 EDTA (Thermo Fisher Scientific) and 1:100 Halt™ protease and phosphatase inhibitor cocktails (Thermo Fisher Scientific) and then pelleted at maximum speed (∼14,000 *g*) for 15 minutes at 4°C. Protein concentrations were quantified using the Pierce® BCA Protein Assay Kit (Thermo Fisher Scientific) following the manufacturer’s instructions. Equal amounts of protein (40 μg) were mixed with 6× Laemmli sample buffer (Bio-Rad) containing 5% β-mercaptoethanol, boiled at 95°C for 5 minutes and were loaded onto Mini-PROTEAN® 4–20% TGX™ stain-free precast gels (Bio-Rad) and separated by SDS-PAGE. Proteins were transferred onto nitrocellulose membranes using the Trans-Blot® Turbo™ Transfer System (Bio-Rad). Membranes were blocked for 1 hour at room temperature in Odyssey® Blocking Buffer (LI-COR) diluted 1:2 in phosphate-buffered saline (PBS), then incubated overnight at 4°C with primary antibodies diluted in Odyssey Blocking Buffer with 0.1% Tween-20 (PBS-T, 1:2). The following antibodies were used for immunoblotting: rabbit anti-HSP90 (Cell Signaling Technologies (CST), 4877, 1:1000), rabbit anti-GAB1 (CST, 3232, 1:1000), rabbit anti-pGAB1(Y627) (CST, 3231, 1:1000), rabbit anti-EGFR (CST, 4267, 1:1000), rabbit anti-pEGFR (Y1068) (CST, 3777, 1:1000), rabbit anti-SHP2 (CST, 3397, 1:1000), rabbit anti-GRB2 (CST, 3972, 1:1000), rabbit anti-NF1 (CST, 14623, 1:1000), rabbit anti-RASA1 (Invitrogen MA4-001, 1:200), rabbit anti-RASAL1 (Abcam, ab168610, 1:1000), rabbit anti-GFP (CST, 2956, 1:1000), rabbit anti-pERK1/2(T202/Y204) (CST, 4370, 1:1000), mouse anti-ERK1/2 (CST, 9107, 1:1000), rabbit anti-pAKT(S473) (CST, 4060, 1:2000), mouse anti-AKT (CST, 2966, 1:2000), rabbit anti-BirA (Agrisera, AS20, 4440, 1:1000), rabbit anti-biotin (CST, 5597, 1:1000), mouse anti-KRAS (Sigma-Aldrich, 3B10-2F2, 1:1000), mouse anti-KRAS (Santa Cruz, sc-30, 1:1000). Primary rabbit antibodies were detected with anti-rabbit IgG (H+L) (DyLight™ 800 4X PEG Conjugate) (CST, 5151, 1:10000). Primary mouse antibodies were detected with anti-mouse IgG (H+L) (DyLight™ 680) (CST, 5470, 1:10000), or HRP-conjugated (Bio-Rad, 1706516, 1:10000) secondary antibodies for fluorescent or chemiluminescent detection (Thermo Scientific, 34076X4), respectively, using the Bio-Rad Chemidoc imaging system. Approximate locations of Kaleidoscope (Bio-Rad) protein size markers (in kDa) are shown in individual blots.

#### Active RAS detection by ELISA and Pull-Down Assays

Active GTP-bound RAS was measured using RAS G-LISA GTPase Activation Assay Kit (Cytoskeleton, Inc.) and the Thermo Scientific™ Active Ras Pull-Down and Detection Kit (Cat. #16117) to detect RAS-GTP levels by ELISA or RAF RBD-based Pull-Down, respectively. Cells were cultured in 10-cm dishes and treated with either vehicle (DMSO) or drugs (GDC-0941, ZSTK474, or 1938), as specified. At about 80% confluency, cells were placed on ice, washed twice with ice-cold PBS, and lysed in 750 μL of ice-cold lysis/binding/wash buffer (provided in the kits) supplemented with Halt™ protease and phosphatase inhibitors. Lysates were incubated on ice for 10 minutes and clarified by centrifugation at 12,000 × g for 15 minutes at 4 °C. The supernatant was collected, and protein concentration was measured using Precision Red Advanced Protein Assay Reagent (in the G-LISA kit) or the Pierce® BCA assay (for Pull-Down assay). For the G-LISA assay, 30 μg per condition (in triplicate) were incubated in the G-LISA plate and the remainder of the assay was performed per manufacturer’s protocol. Absorbance at 490 nm (background subtracted) was measured using a BioTek Synergy plate reader to measure RAS-GTP signal. Positive control was provided in the kit, and negative control was buffer only without protein. For the Active RAS Pull-Down assay, 1000 μg total protein for each condition was incubated with 80 μg of Raf1-RBD agarose (provided in the kit) for 1 hour at 4°C with gentle rotation to selectively isolate GTP-bound RAS. Following incubation, beads were pelleted by centrifugation at 6,000 × g for 30 seconds and washed three times with lysis/binding/wash buffer. The final pellet was resuspended in 2× SDS-PAGE sample buffer, boiled at 95°C for 5 minutes, and subjected to SDS-PAGE followed by immunoblotting using the KRAS monoclonal antibody (Santa Cruz, sc-30, 1:1000). 10% of each lysate was reserved and analyzed in parallel to assess total RAS levels.

#### Immunoprecipitation

To isolate KRAS or GAB1 and their associated protein complexes, GFP-tagged KRAS^WT^ and/or mCherry-tagged GAB1 constructs were stably expressed in 8988T-KO cells. Cells were cultured in 10-cm dishes and harvested when reaching ∼80% confluency. Where indicated, cells were treated with small-molecule drugs (*e.g.,* GDC-0941, 1938) or vehicle (DMSO) for 0.5 hours prior to lysis. Cells were washed twice with cold PBS and lysed on ice for 30 minutes in freshly prepared lysis buffer (50 mM Tris-HCl pH 7.5, 150 mM NaCl, 1% NP-40, 1 mM EDTA, 10% glycerol) supplemented with Halt™ protease and phosphatase inhibitors. Lysates were clarified by centrifugation at 20,000 × g for 15 minutes at 4°C, and the protein concentration was determined using the BCA assay. For immunoprecipitation, 25 μL of GFP-Trap or RFP-Trap Magnetic Agarose beads (Chromotek) per sample were equilibrated according to the manufacturer’s instructions and incubated with clarified lysates (1 mg total protein) for 1 hour at 4°C with gentle rotation. Beads were then washed three times with wash buffer to remove non-specifically bound proteins. Bound proteins were eluted in 2× SDS-PAGE sample buffer by heating at 95°C for 10 minutes and subjected to immunoblotting.

#### Live-cell confocal microscopy

Live-cell imaging was performed using an Andor Benchtop Confocal microscope (BC43) equipped with a 60× oil immersion objective (NA 1.4). Prior to imaging, the system was equilibrated for 30 minutes to ensure stable temperature and humidity. Cells were cultured in glass-bottom dishes and maintained during imaging in an incubation chamber set to 37°C with 5% CO_2_. For ERK-KTR imaging to monitor ERK activity dynamics, either GDC-0941 (2 μM) or vehicle control (DMSO) was added at the 5-minute timepoint, and time-lapse imaging was continued for an additional 35 minutes at 30-second intervals. For imaging analysis of GAB1 and KRAS co-localization, cells were first treated with DMSO for 3 minutes, followed by sequential addition of 1938 (1 μM) and GDC-0941 (2 μM). Time-lapse imaging was performed over a total duration of approximately 30 minutes at 15-second intervals.

#### Proximity protein labeling and streptavidin pull-down of biotinylated proteins

Proximity protein labeling was performed using cells expressing either BirA-tagged KRAS constructs (WT, G12V, C185S, HVR) or MiniTurbo-KRAS^WT^ constructs. Cells were seeded in 15-cm dishes and cultured until approximately 80% confluency. For BirA-based labeling, cells were incubated for 16 hours in DMEM supplemented with 50 μM biotin and either 2 μM GDC-0941 or DMSO as a control. For MiniTurbo-based labeling, cells were treated with biotin and indicated drug conditions (DMSO, GDC-0941, 1938, or their combination) for 4 hours. A no-biotin control was included to assess background signal from endogenously biotinylated proteins. After labeling, cells were washed five times with ice-cold PBS to remove residual biotin, then harvested by scraping. Cell pellets were lysed in ice-cold RIPA buffer (50 mM Tris-HCl pH 7.5, 150 mM NaCl, 1% NP-40, 1 mM EDTA, 1 mM EGTA, 0.1% SDS, 0.5% sodium deoxycholate, and protease inhibitors) and incubated on a nutator for 1 hour at 4°C. Lysates were centrifuged at maximum speed (≥20,000 × g) for 30 minutes at 4°C, and the cleared supernatants were incubated overnight with pre-washed streptavidin-conjugated Dynabeads (Thermo Fisher) at 4°C with gentle rotation. Beads were washed sequentially with RIPA buffer, once with 2% SDS in 50 mM Tris-HCl (pH 7.4), and three times with 100 mM ammonium bicarbonate (pH 8.0). For on-bead digestion, proteins were reduced with 10 mM dithiothreitol (DTT) for 30 minutes at 50°C and alkylated with 100 mM iodoacetamide for 30 minutes at room temperature in the dark. Proteins were then digested overnight at 37°C with 1 μg of trypsin in 100 mM ammonium bicarbonate. Eluted peptides were dried via vacuum centrifugation and resuspended in buffer containing 98% HPLC-grade water, 2% acetonitrile (ACN), and 0.1% formic acid for LC-MS/MS analysis.

#### Mass Spectrometry and quantitative proteomics

All the samples were measured by data-independent acquisition mass spectrometry (DIA-MS), as described previously^75,76^, using an Orbitrap Fusion Tribrid mass spectrometer (Thermo Fisher Scientific) instrument coupled to a nanoelectrospray ion source (NanoFlex, Thermo Fisher Scientific) and EASY-nLC 1200 systems (Thermo Fisher Scientific). A 120-min gradient was used for data acquisition at a flow rate of 300 nL/min with the temperature controlled at 60°C using a column oven (PRSO-V1, Sonation GmbH, Biberach, Germany). The DIA-MS method consisted of one MS1 scan and 33 MS2 scans of variable isolated windows with 1 m/z overlapping between windows. The MS1 scan range was 350 – 1650 m/z, and the MS1 resolution was 120,000 at m/z 200. The MS1 full scan AGC target value was set to be 2e6 and the maximum injection time was 100 ms. The MS2 resolution was set to be 30,000 at m/z 200 with the MS2 scan range of 200 – 1800 m/z and the normalized HCD collision energy at 28%. The MS2 AGC was set to be 1.5e6 and the maximum injection time was 50 ms. The default peptide charge state was set to 2. Both MS1 and MS2 spectra were recorded in profile mode. DIA-MS data analysis was performed using Spectronaut v16-19^77^ with the directDIA algorithm by searching against the Human SwissProt fasta file^78,79^. Both peptide and protein FDR cutoffs (q-value) were controlled below 1%. All other settings in Spectronaut were kept as Default.

#### CRISPR screens

Genome-scale CRISPR screening was performed using the Brunello sgRNA knockout library^46^ in independent pools of 8988T (KRAS^G12V^, *KRAS*-intact) and *KRAS*-deficient (*KRAS*-KO) clones (4 clones per pool). High-titer lentiviral supernatant was obtained from the Genetics Perturbation Platform (Broad Institute) and applied to each cell pool with a target multiplicity of infection (MOI) of 0.3 with a target sgRNA representation of 500x. ∼150 million cells were transduced by spinfection in 12-well plates (1.5 million cells per well) and treated with puromycin 2 µg/mL for 7 days starting one day post-transduction. Cells were passaged at ∼80% confluency and sufficient cells were passaged to always maintain 500x representation of the sgRNA library. Cell pellets were collected and flash frozen at three time points post-transduction for each condition: Day 4 (D4), serving as the initial time point and representing the starting library distribution; Day 8 (D8), an early time point at the conclusion of puromycin selection; and a final time point corresponding to 14 population doublings post-transduction (Day 22 for *KRAS*-intact cells and Day 32 for *KRAS*-KO cells). Genomic DNA was isolated from cell pellets using a Qiagen Midi kit per manufacturer’s protocol. sgRNA sequences were PCR amplified with barcodes and Illumina sequencing adaptors using ExTaq DNA polymerase (Clontech), purified with Agencourt AMPure XP SPRI beads (Beckman Coulter), and sequenced on Illumina NextSeq to obtain 100-bp reads by the MIT BioMicroCenter.

#### *In vitro* drug assays

Cells were seeded in 96-well plates at a density of 1000 cells per well in complete growth medium and allowed to adhere overnight. The following day (designated Day 0), cells were treated with vehicle (DMSO) or indicated drugs at specified concentrations. On Day 3, cell viability was assessed using the CellTiter-Glo® Luminescent Cell Viability Assay (Promega) according to the manufacturer’s instructions. Luminescence was measured using BioTek Synergy plate reader and normalized to DMSO. Each well in a single experiment was treated as a technical replicate, and data presented are representative of at least two independent experiments.

### QUANTIFICATION AND STATISTICAL ANALYSIS

#### Immunoblot quantification

Protein quantification from western blots was performed using the Fiji gel analyzer, and data for each biologic replicate along with mean ± SD are presented in bar graphs, as described in the figure legends. Statistical analyses were performed using Prism (GraphPad). Comparisons between biological replicates of two groups (e.g., drug vs. solvent) were performed using paired two-tailed t-test. Multiple group comparisons were performed using one-way ANOVA or repeated measures ANOVA (rmANOVA) followed by post-hoc comparisons using Dunnet’s test or Šidák’s test for multiple comparisons, as described in the figure legends. *p* < 0.05 was used as level of significance for all statistical analyses.

#### Quantification of ERK activity using ERK-KTR

ERK activity was quantified using the ERK-Kinase Translocation Reporter (ERK-KTR) by measuring the relative cytoplasmic-to-nuclear fluorescence intensity. Cells expressing ERK-KTR were imaged by fluorescence confocal microscopy under identical acquisition settings. Image analysis was performed using Fiji. For each cell, regions of interest (ROIs) corresponding to the whole cell and nucleus were manually drawn based on morphology or co-staining with DAPI. Mean fluorescence intensities were measured for both the whole cell and nucleus ROIs. ERK activity was calculated as the ratio of cytoplasmic to nuclear fluorescence intensity at each time point using the following formula: ERK Activity = (Whole Cell Intensity – Nuclear Intensity) / Nuclear Intensity. To enable comparison across individual cells, ERK activity values were normalized to the mean activity of the baseline condition (DMSO-treated) for each cell.

#### Quantification of GAB1 and KRAS membrane recruitment

To quantify the membrane recruitment of GAB1 and KRAS, cells expressing fluorescently tagged constructs were imaged using confocal microscopy under standardized acquisition settings. Image analysis was conducted in Fiji. For each cell, five line profiles were manually drawn perpendicular across the plasma membrane within defined regions of interest (ROIs). Along each line, the membrane intensity was measured at the peak corresponding to the plasma membrane (PM) signal, while the adjacent cytosolic intensity was measured six pixels inward from the membrane on the cytoplasmic side, referred to as the adjacent cytosol. The membrane recruitment ratio for each line was calculated, which corresponded to the PM/cytosol intensity ratio. The average of the five line-derived ratios was taken as the value for each cell. At least 50 cells were analyzed across 10 independent experiments. Background fluorescence was subtracted, and data were presented as mean ± SD and statistical comparisons were made using one-way ANOVA with Šidák post-hoc test for multiple comparisons. Pearson correlation analysis was performed to determine associations between two continuous variables. *p* < 0.05 was used as level of significance for all statistical analyses.

#### MAGeCK analysis of CRISPR screening results

Raw sequencing reads from each sample were processed using the MAGeCK (Model-based Analysis of Genome-wide CRISPR-Cas9 Knockout) software suite. The “mageck count” function was used with default parameters to quantify sgRNA read counts across all samples, generating a count matrix for downstream analysis. Differential sgRNA and gene-level enrichment analyses were performed using the “mageck test” function. Read counts were normalized using the non-targeting control sgRNAs included in the Brunello library to control for global shifts in sgRNA abundance and ensure robust estimation of enrichment or depletion signals. Comparisons were made between the final time point (14 population doublings) vs. D4 for each condition independently. MAGeCK computed log_2_ fold changes (LFC), p-values, and false discovery rates (FDRs) for both negative and positive selection at the sgRNA and gene levels, based on differences in sgRNA abundance relative to the initial D4 library representation. The resulting enrichment statistics were used to identify genes with significant dropout or enrichment over time, revealing putative context-dependent vulnerabilities in *KRAS*-intact versus *KRAS*-KO backgrounds.

#### Proteomics data analysis and visualization

Quantitative proteomics data were analyzed using R (v4.3.0). Protein intensities were log_2_-transformed, and differential abundance between DMSO- and drug-treated samples was assessed for each genotype (WT, G12V, C185S, HVR) using the limma package (v3.56.2) with empirical Bayes moderation. FDRs were computed using the Benjamini–Hochberg method. A curated list of RAS-pathway genes, annotated by category (*e.g.,* Core RAS Family, Effectors, GAPs, Scaffold/Adaptors, RTKs), was used to generate heatmaps and annotate volcano plots. Heatmaps were created using pheatmap (v1.0.12) after row-wise z-score normalization, with samples displayed unclustered and genes grouped by category. Volcano plots were generated using ggplot2 (v3.4.2) and ggrepel (v0.9.3), with a fixed y-axis scale across conditions.

